# Microfluidic sieve and detector for rapid ultrasensitive assays with single-molecule sensitivity

**DOI:** 10.1101/2025.10.30.685461

**Authors:** Geunyong Kim, Houda Shafique, Molly L. Shen, Andy Ng, David Juncker

**Affiliations:** Biomedical Engineering Department, McGill University, Montreal, QC, Canada; Victor Phillip Dahdaleh Institute of Genomic Medicine, McGill University, Montreal, QC, Canada

## Abstract

The advent of ultrasensitive assays enabled many new research and clinical applications, but the time-to-result is hours and has not significantly improved over time, limiting their potential. We introduce the microfluidic sieve and detector (MFSD) that can sieve 200 microlitres in just one minute, and quantify captured analytes in four minutes, down to zeptomolar (10^-19^M) concentration. The MFSD comprises half a million 8-μm-diameter pores that bind analytes upon Brownian motion-induced wall collision. Conventional ensemble models and a stochastic Brownian agent dynamics introduced here reveal rapid, predictable and tuneable analyte capture under flow. Digital sandwich assays are completed by sequentially flowing the sample and reagents, partitioning the pores, and digitally revealing single ‘trapped’ analytes by enzymatic amplification. The MFSD tests are specific and reproducible, exhibit a large dynamic range, and are simple, tuneable and versatile, enabling measurement of different proteins including influenza A nucleoprotein in clinical samples, and even nucleic acids while being both faster and more sensitive than PCR. As such, the MFSD could open a new chapter for analysis and diagnostics.

## Main

Rapid testing offers many advantages for diagnostics, but common approaches face trade-offs between sensitivity, time-to-result, simplicity and cost. Both protein and nucleic acid tests rely on binding between an analyte and a capture probe following their encounter upon diffusive, and sometimes convective, transport. Microfluidics has notably been used to shorten time-to-result by miniaturizing containers to speed-up diffusive mass transport^1–3^; however, this comes at the expense of sensitivity because the inclusion of a statistically representative number of analytes requires ever larger sample volumes, while microchannels only accommodate microscopic volumes. A strategy to both accommodate larger volumes yet minimize the time and distance of diffusive transport between analyte and probes is to disperse the latter in solution, either directly or attached to microparticles^4,5^. Nucleic acid tests^6–11^ disperse probes and use complementary base pairing recognition to initiate exponential amplification (*e.g.* polymerase chain reaction and isothermal amplification) or collateral cleavage of reporter oligonucleotides in CRISPR-based tests, which enable ultrasensitive detection, but at a comparatively long time-to-result of tens of minutes for low concentrations, while being limited by a small volumetric capacity of typically a few microliters only. Protein immunoassays have historically been less sensitive and conventional enzyme-linked immunosorbent assays (ELISAs) only use linear amplification^12^ and are lengthy. The law of mass action and Langmuir isotherms suggest that as higher sensitivity is sought, the finite affinity of reagents will limit performance, and the time required to complete the assay will increase^13,14^. A breakthrough in ultrasensitive assays was achieved by digital counting of single binding events. The most common strategy relies on dispersing large numbers of antibody-coated microparticles in the sample, thus shifting the binding equilibrium towards higher antibody-analyte binding, followed by incubation with detection antibody-enzyme conjugate for signal generation, and partitioning into ‘dead-end’ microwells or droplets, enabling digital readout^14–24^. However, the protocol remains lengthy (hours), and the instrumentation complex. Here we describe a microfluidic sieve and detector (MFSD) that can sieve molecules very quickly using an isoporous membrane with each open-ended, micrometer-scale pore forming a Brownian affinity trap (BAT). Under commensurate sample flow, Brownian-motion induced wall collision leads to analyte capture within the BATs, which upon partitioning, turn into a digital detector array for quick counting and quantification of every ‘trapped’ analyte.

## Results

The MFSD test is completed by sieving the sample for 1 min, capturing analytes in BATs, and by detecting the captured analytes in 4 min via sandwich binding, then partitioning of each BAT into a closed compartment by oil capping and enzymatic conversion of a soluble substrate in each BAT, all within only 5 min, and achieves zM–aM–fM ultrahigh sensitivity and 6 OM dynamic ranges with good reproducibility of ≲15% coefficient of variation (CV), for both protein and nucleic acid analytes (Fig. 1a and see Extended Data Fig. 1 for large-area MFSD images with digital signals). In this study, the MFSD comprised 500,000 BATs at a density of 800,000 pores cm^-2^ (40% open ratio) to permit rapid sieving of a sample at a high overall flow rate while meeting the commensurate, slow flow condition for a BAT, each of which is a cylindrical, open-ended micropore with a radius *R*_BAT =_ 4 µm and a length *L*_BAT =_ 20 µm (Fig. 1b,c). Akin to other digital assays, a single enzyme rapidly generates easily detectable fluorescence in the microscopic 1-pL BAT, thus permitting binary classification of BATs into positive (‘1’) and negative (‘0’). As concentration increases, more BATs turn positive while the probability of more than one analyte per BAT also increases, which is predicted and compensated for by Poisson statistics and by calculating the average number of enzymes per BAT (AEB). For high percentage of positives (≳50%), AEB calculations become imprecise, but conversely analogue readout becomes possible simply by measuring the average fluorescence signal of the MFSD (see method for signal readout details). By combining digital (MFSD-D) and analogue (MFSD-A) detection, the dynamic range of an MFSD test can be extended^21^. Three critical features for MFSD tests are (i) arrays of open-ended micropores (*i.e.* BATs) that are ∼1,000 times larger than the sieved ‘nanoparticles’, facilitating clogging-free flow and affinity sieving, while minimizing the required pressure head, (ii) exclusive capture of analytes inside of the BATs, and not on the top and bottom surfaces between the BATs to preserve proportionality between number of molecules and positive counts, and (iii) leak-free capping with oil to partition BATs and enable digital readout.

**Fig. 1.**
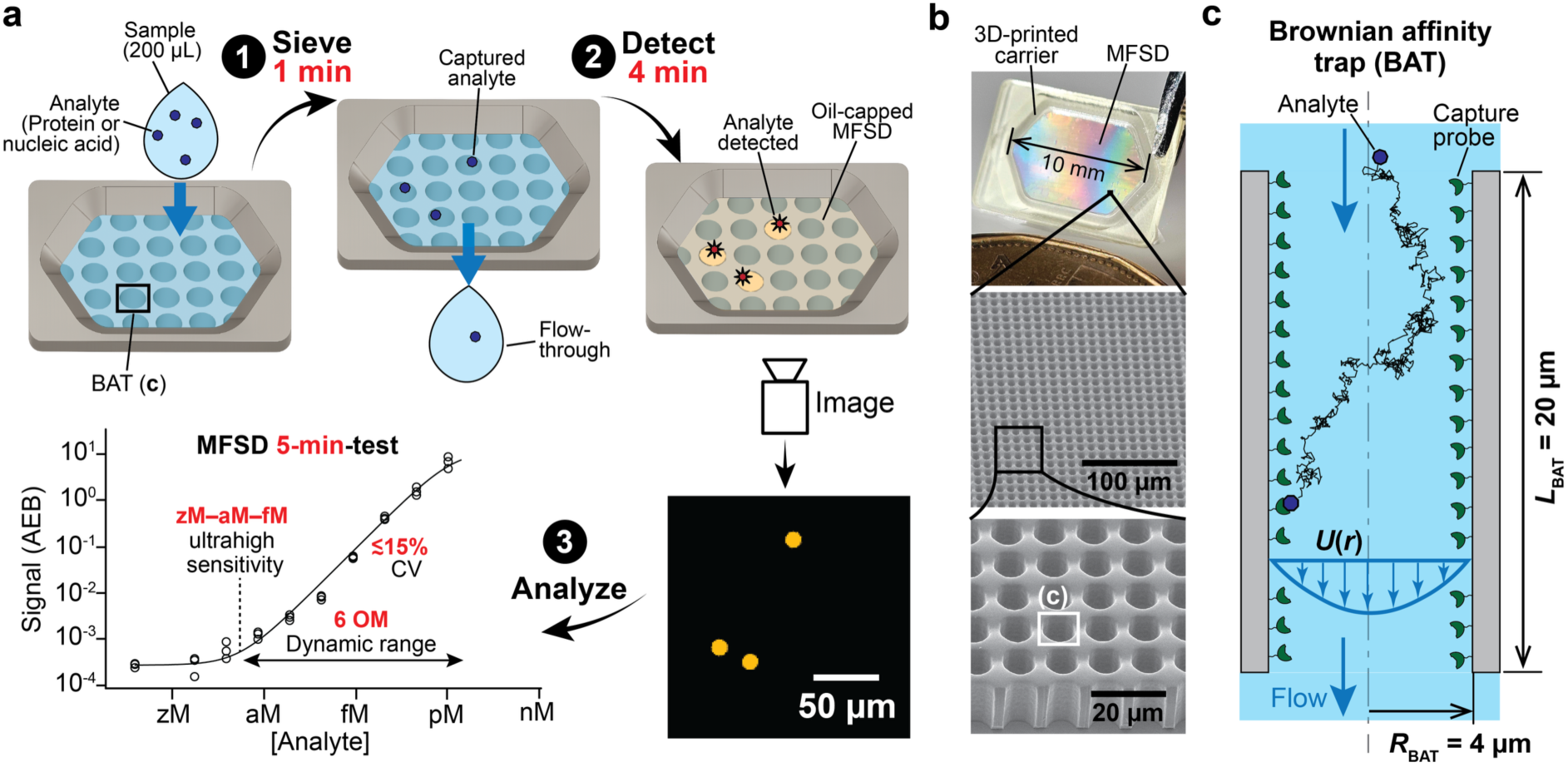
Microfluidic sieve and detector (MFSD) made from an array of Brownian affinity traps (BATs) captures and detects single analytes in 5 min. **a**, Conceptual MFSD test procedure. The sample is sieved, reagents and buffers are sequentially flowed through the MFSD, BATs are partitioned by oil capping, and BATs that contain one or more captured analytes become fluorescent upon enzymatic amplification, digitally revealing single molecule binding and realizing a rapid, ultrasensitive test. b, MFSD with 500,000 BATs bonded to a 3D-printed carrier with successive magnification. c, Cross-section view of a 4 μm radius and 20 μm long BAT. A BAT is a microchannel coated with capture probe under commensurate flow promoting Brownian motion-induced wall collision of an analyte and affinity binding by the immobilized capture probe. The black trace reveals the path of a captured analyte.

### Theoretical capture efficiency of the MFSD and the BAT

A key metric of the MFSD test is the analyte capture efficiency *η* of both the MFSD and the BAT defined as the fraction of trapped to total analytes flowed through the MFSD and BAT. We calculate *η* theoretically by a (i) stochastic Brownian agent dynamics (SBAD) method we introduce, and by ensemble (ii) analytical and (iii) finite element method (FEM) calculations of binding reactions. SBAD is a computational simulation of the Brownian motion of an analyte (the Brownian agent) as a random walk in the BAT under flow, while wall collision and affinity binding are simulated as stochastic events^25–27^. SBAD visualizes single analytes, is intuitive and accounts for the stochasticity and stochastic variance that are ignored by ensemble models, while accurate *η* are obtained by Monte Carlo estimation upon repeated simulations. We presume geometrical ergodicity implying that the *η* of a single BAT calculated by SBAD is equivalent to the *η* of a MFSD, and concentration ergodicity implying that ensemble calculations of the reaction *η* of a BAT as the concentration change from inlet to outlet is equivalent to the capture *η* of single molecules flowing through a BAT. For all models, we consider that analytes enter the BAT pore with a length *L*_BAT a_nd a radius *R*_BAT,_ at an average flow velocity *Ū* following a parabolic Poiseuille flow profile and are subject to Brownian motion according to the diffusion coefficient *D* [m^2^ s^-1^], see Fig. 1c.

For the ensemble models, we consider the on-rate *k*_on [_m^3^ mol^-1^ s^-1^] under first-order Langmuir kinetics, the surface density of the immobilized capture probe *Γ* [mol m^-2^], and the surface-to-volume ratio 2/*R*_BAT. W_e define *κ*_on =_ 2*k*_on*Γ*_/*R*_BAT [_s^-1^] as the effective on-rate and *τ*_res =_ *L*_BAT/_*Ū* [s] as the average residence time of the sample. In this study, we operate under conditions of large *κ*_on (_∼10^-1^–10^2^ s^-1^), low density of bound analyte, and short timescales with low off-rates *k*_off (_∼10^-6^ s^-1^), so that dissociation can be neglected in the analysis. The Péclet (Pe) and Graetz (Gz) numbers quantify the ratio of convective to diffusive transport and are > 1 for all experiments, indicating convection-dominated transport. Damköhler numbers (Da) express the ratio of the reaction to mass transport rates and we consider the first Damköhler number, Da_I =_ *κ*_on*τ*res r_epresenting the ratio of reaction rate to convective mass transport and a surface Damköhler number, Da_s =_ *k*_on*Γ*_/*k*_c w_ith *k*_c [_m s^-1^] being the mass transfer coefficient used in thin film theory, which is the ratio of reaction rate to the total mass transport to the surface including both convection and diffusion (Fig. S1 and Supplementary Information)^1,28,29^. We calculate the ensemble reaction efficiency *η* of a BAT following classical derivations^29–31^ and find (Fig. S2 and Supplementary Information for the detailed derivation):

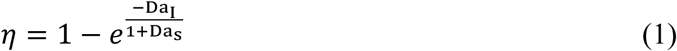

For most experiments in this study, Da_s <_ 1, hence we obtain:

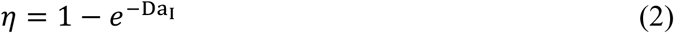

Hence, *η* is adjustable by varying *κ*_on (_*e.g.* via the surface density of capture probes) and *τ*_res v_ia *Ū*. The FEM ensemble calculations directly solve the coupled convection-diffusion-reaction equations without simplifying assumptions and provide spatially resolved concentration fields, from which *η* is obtained as the ratio of the outlet to inlet average concentration, Fig. 2a,b.

**Fig. 2.**
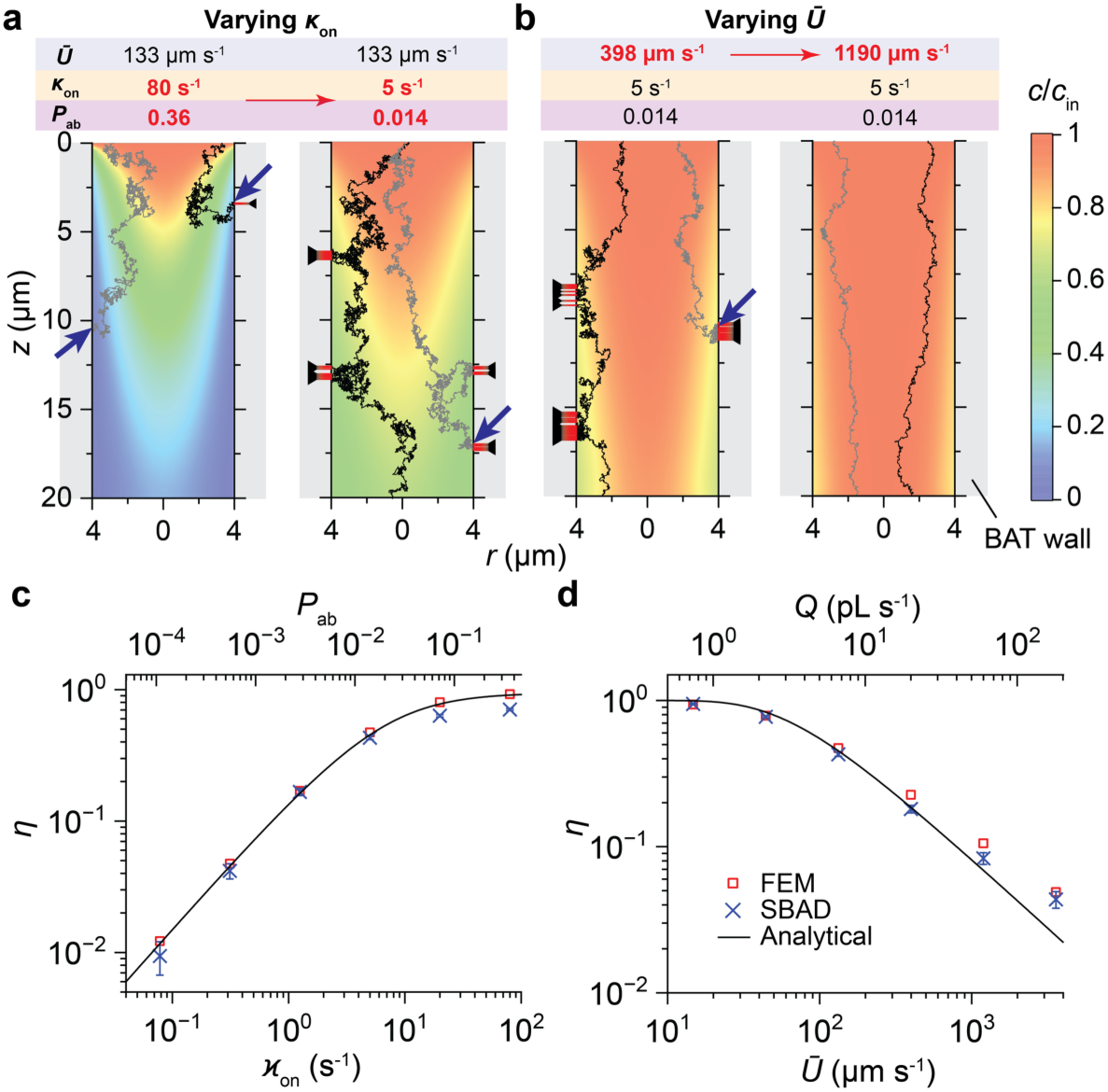
Stochastic Brownian agent dynamics (SBAD) traces and analytical, SBAD and FEM calculations of BAT capture efficiency *η* for different conditions. **a**, SBAD example traces overlaid on FEM simulation for a model 100 kDa globular protein analyte with a diffusion coefficient *D* = 80 µm^2^ s^-1^ with varying effective on-rate *κ*on and affinity binding probability *P*ab at a fixed average flow speed, *Ū* = 133 µm s^-1^, and with **b**, varying *Ū* at fixed *κ*on = 5 s^-1^ and *P*ab = 0.014. Arrows record analyte binding while red bars with a black arrow tail indicate wall collision without binding. The colour bar indicates the ratio, *c*/*c*in, where *c* is the local analyte concentration and *c*in is the inlet concentration. **c**, *η* as a function of *κ*on at *Ū* = 133 µm s^-1^ and **d**, of *Ū* at *κ*on = 5 s^-1^. SBAD error bars represent the two-sided 95% confidence interval of SBAD with *N* = 5,000 and increase for low *η* due to stochastic noise.

SBAD analysis is described in detail in Supplementary Information including the computational algorithm in Fig. S3. Example traces visualizing random-walk analyte trajectories under different flow conditions and varying surface affinity, are shown as slow-motion, time-stamped animations in Supplementary Videos S1–S3, and as traces superposed with FEM ensemble models of concentration in Fig. 2a,b. To calibrate SBAD, we introduce the affinity binding probability upon wall collision (*P*_ab)_ as a free parameter. Benchmarking against *η* obtained from the FEM calculations, calibration was performed for the model system with a 100 kDa globular protein analyte with *D* = 80 µm^2^ s^-1^ flowing at *Ū* = 10 µm s^-1^. *P*_ab w_as determined and found to be proportional to *κ*_on (_Extended Data Fig. 2 and Supplementary Information) for a wide range of *κ*_on,_ and constant relative to *τ*_res (_and *Ū*) across a broad range, Fig. 2c,d. A lower *κ*_on (_and lower *P*_ab)_ result in a higher number of non-binding collisions, and a lower *τ*_res i_n fewer wall collisions, both contributing to lower *η*. The calibrated SBAD was consistent with both FEM and analytical calculations while capturing stochastic variances of single-analyte binding in BATs.

### MFSD membrane manufacturing and functionalization

MFSDs were made by moulding and photopolymerization of a perfluoropolymer forming an isoporous membrane almost identical to the one used for size-exclusion-based filtration of circulating tumour cells^32,33^, but used here for Brownian affinity trapping of molecular analytes inside of the pores. A crucial prerequisite for MFSD assays is two-sided hydrophobicity of the membrane to both prevent unwanted binding of the analytes on the ‘top surface’, outside of the BAT, which could confound assay results, and to form a water-tight seal upon capping with fluorinated oil and partition each BAT for digital assay readout. We developed float-flip-float process to exclusively functionalize the inside of BATs with capture probes and preserve hydrophobicity on the top and bottom surfaces of MFSDs as described in Supplementary Information and shown in Extended Data Fig. 3.

### Experimental efficiency of the MFSD and of the BAT

We systematically evaluated MFSD and BAT sieving *η* experimentally with an array of 500,000 BATs, using the experimental setup shown in Fig. 3a. The captured analytes were directly detected via a one-step MFSD test, Fig. 3b. Briefly, MFSDs were functionalized with biotin as a capture probe, and streptavidin-β-galactosidase conjugate (SβG) was used as a combined “analyte-detection probe-enzyme”. The direct detection of the captured analytes eliminates confounding factors arising from downstream steps such as detection probe and enzyme incubations that contribute to false positives and negatives. We independently established *κ*_on =_ 7.44 s⁻¹ and *P*_ab =_ 0.035 with high density functionalization (for details on *κ*on, see Extended Data Table 1, and Supplementary Figs. S4 and S5).

**Fig. 3.**
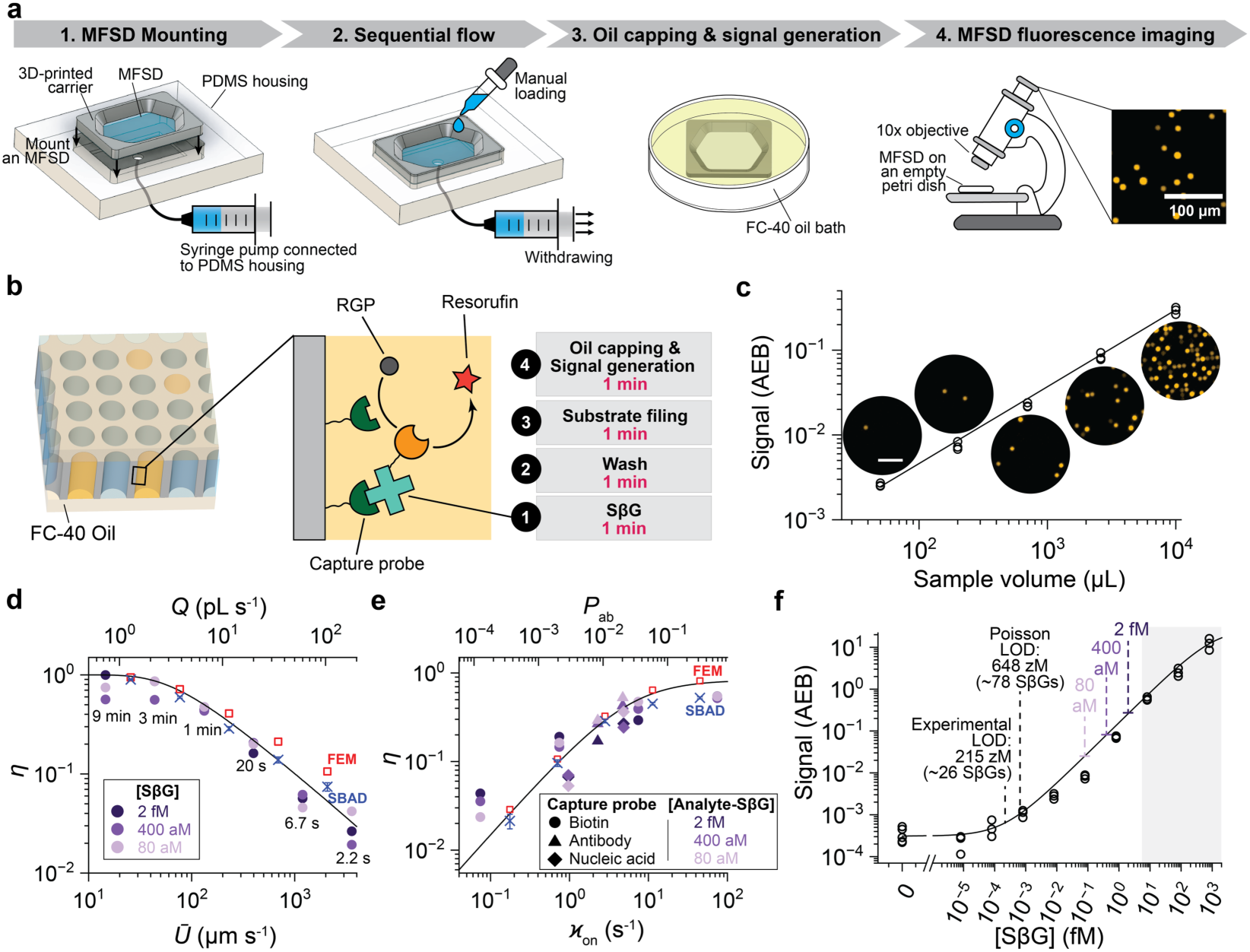
Experimental one-step MFSD test sieving efficiency *η* is concordant with theory. MFSD tests were conducted using a 200 µL solution containing streptavidin-β-galactosidase (SβG, MW = 530 kDa and *D* = 45.9 µm^2^ s^-1^) flowed through an array of 500,000 BATs in 1 min with *Ū* = 133 µm s^-1^ and with *κ*on = 7.44 s^-1^, unless indicated otherwise. **a**, MFSD test experimental setup. Liquids are manually loaded and sequentially flowed through the MFSD mounted on a PDMS housing by a syringe pump, followed by signal generation in an oil bath and fluorescence microscope imaging. **b**, Workflow of one-step MFSD test with SβG “analyte-detection probe-enzyme”. **c**, The number of captured analytes (i.e., AEB) as function of sample volume follows a linear relationship with a slope of 0.91 and *R*^2^ = 0.99. Insets are illustrative MFSD images. Scale bar = 50 µm. **d**, Experimental sieving efficiency *η* as a function of flow speed *Ū* (and corresponding flow rate *Q* and incubation times) and **e**, of *κ*on (and corresponding *P*ab) for different capture probes (biotin, antibody, nucleic acid each with its corresponding analyte-SβG system) and varying surface density. *η* was measured for analyte concentration of 2 fM, 400 aM and 80 aM, and found to be invariable as expected. Line is analytical model, squares are FEM and ‘×’ is SBAD. **f**, MFSD standard binding curve for this 4-min test (see **b**). The white and greyed areas of the curve correspond to MFSD-D and MFSD-A, respectively. The experimental limit of detection (LOD) was defined as the concentration yielding an AEB equal to the mean plus three standard deviations of blank replicates. The Poisson LOD was independently estimated from three times the Poisson standard deviation of the mean background count, the measured capture efficiency and the sample volume of 200 µL.

In typical experiments, 200-µL-samples were sieved at a flow velocity of *Ū* = 133 µm s^-1^ for 1 min and the captured molecules were detected following sequential 1-min steps of washing and flowing of resorufin-β-D-galactopyranoside (RGP), followed by oil capping and enzymatic conversion of RGP into fluorescent resorufin. We evaluated the number of captured molecules as a function of the volume over 2 OM and observed the expected proportionality, Fig. 3c. The MFSD and BAT efficiency *η* was quantified by calculating the ratio of captured to total number of molecules flowed under varying conditions, and compared to theory. We varied flow velocity (and corresponding flow rate) and *κ*_on (_and corresponding *P*_ab)_ over at least 2 OM each. *κ*_on w_as modulated by varying the surface density of biotin serving as the capture probe by ratiometric incorporation of spacer molecules and via different capture probe types. We compared experimental results with analytical, FEM and SBAD predictions, and found strong concordance in all cases, Fig. 3d,e, thus mutually validating and reinforcing theory (notably SBAD analysis) and experimental findings. The data showcase rapid, predictable and tuneable analyte capture by microfluidic sieving of 200 μL in just 2 s with *Ū* = 3,580 µm s^-1^ for *η* ∼ 2% (characteristic time scales of convection ∼10^-3^ s << reaction and diffusion ∼10^-1^ s), and in 1 min with *Ū* = 133 µm s^-1^ for *η* ∼ 50% (with comparable characteristic time scales across convection, reaction and diffusion, ∼10^-1^ s), for diverse capture probes including biotin, antibodies and nucleic acids, and with linearly increasing numbers with volume.

Next we evaluated how a high *η* ∼ 50% could benefit assay performance for the one-step test by establishing a standard binding curve for varying concentrations of SβG. We used dynamic thresholding to distinguish positive and negative BATs that proved robust with a coefficient of variation (CV) of 29%, but with residual false-positives of 3.4 × 10^-4^ AEB on average at zero concentration (∼170 classified positive BATs among 500,000 BATs), Fig. 3f. Both the stochastic analyte capture with its standard error (that can be evaluated using the SBAD simulation), and stochastic sampling with Poisson noise, each leading to increased fluctuations scaling with the square root, contribute to CV, and become significant for low analyte numbers. We set the experimental signal threshold at three standard deviations above the blank mean, yielding a threshold of 6.4 × 10^-4^ AEB. Separately, the Poisson noise of the false-positive count (3×√170 = 39) was used to consider sampling stochasticity at zM concentrations where only a few molecules are present in the 200 µL sample (See Supplementary Information for details). The overall noise fluctuated from 42% below the limit of detection (LOD) to 11% in the linear range, which we consider to be good for a manual test. The readout accuracy drops for AEB > 0.5 because increasing number of BATs will have more than one analyte, and for test results above this threshold average fluorescence signals are quantified by MFSD-A readout. With the combined MFSD-D and MFSD-A (Extended Data Fig. 4), we obtained experimental and Poisson LODs of 215 zM and 648 zM, corresponding to expected 26 and 78 molecules in a 200 µL, respectively, both within the zM range, together with a dynamic range of ∼6 OM in a 4-min test, which is remarkable.

### Sandwich MFSD test for protein quantification and influenza A virus detection from nasopharyngeal swab samples

We extended MFSD tests to sandwich assays for proteins with a time-to-result of 5 min accounting for capture and digital detection of molecules. Similar to a conventional ELISA, the sandwich MFSD test is based on sequentially introducing the sample containing the analyte, biotinylated detection antibody and the streptavidin-enzyme conjugate (*i.e.* SβG in our case), leading to an enzymatic sandwich complex, which then produces fluorescent resorufin from RGP (see Fig. 4a,b). BATs were functionalized with capture antibodies against IL-4 or influenza A nucleoprotein. For a 200 µL sample flowed through the MFSD in 1 min at 133 µm s^-1^, Eqs. 1 and 2 predict *η* of 96.2% and 3.67%, respectively, based on their diffusion coefficients and literature-reported *k*_on v_alues^34,35^ (See Table S1 for detailed calculation of *η*). The sandwich format introduces additional sources of noise, and most significantly the non-specific binding of either the detection probe or of SβG each contribute to false positive BATs. False positives quickly degrade assay sensitivity and performance, and should be kept low^22^. We thus systematically optimized blocking, reagent concentration, flow time, flow speed to balance conversion of capture analytes into positive BATs, non-specific binding and assay times. With 10 nM detection antibody and 4 pM SβG each flowed for 1 min at 66.3 µm s^-1^, the background signal for negative controls was ∼2 × 10^-3^ AEB, *i.e.* ∼1,000 positive BATs for an MFSD with 500,000 BATs. Standard curves for IL-4 and influenza A nucleoprotein were generated with the 5-min MFSD test and with a classical 4-h ELISA in microplates, Fig. 4c,d. The LODs of the ELISA were 118 fM (1.77 pg ml^-1^) and 918 fM (52.1 pg ml^-1^), respectively, and of the MFSD 325 aM (4.88 fg ml^-1^) and 2.00 fM (113 fg ml^-1^), respectively. The dynamic ranges were ∼ 2 OM for the ELISA, and 6 OM for the combined MFSD-D and MFSD-A. The MFSD was thus 50 times faster and ∼350 times more sensitive, and dynamic range was extended not only at the low, but also at the high concentration range by 2–3 OM *vs* ELISA. The LOD and CVs of the MFSD test are similar or better compared to other ultrasensitive assays, but with a time-to-result of only 5 min *vs* 2–4 h^20–24^.

**Fig. 4.**
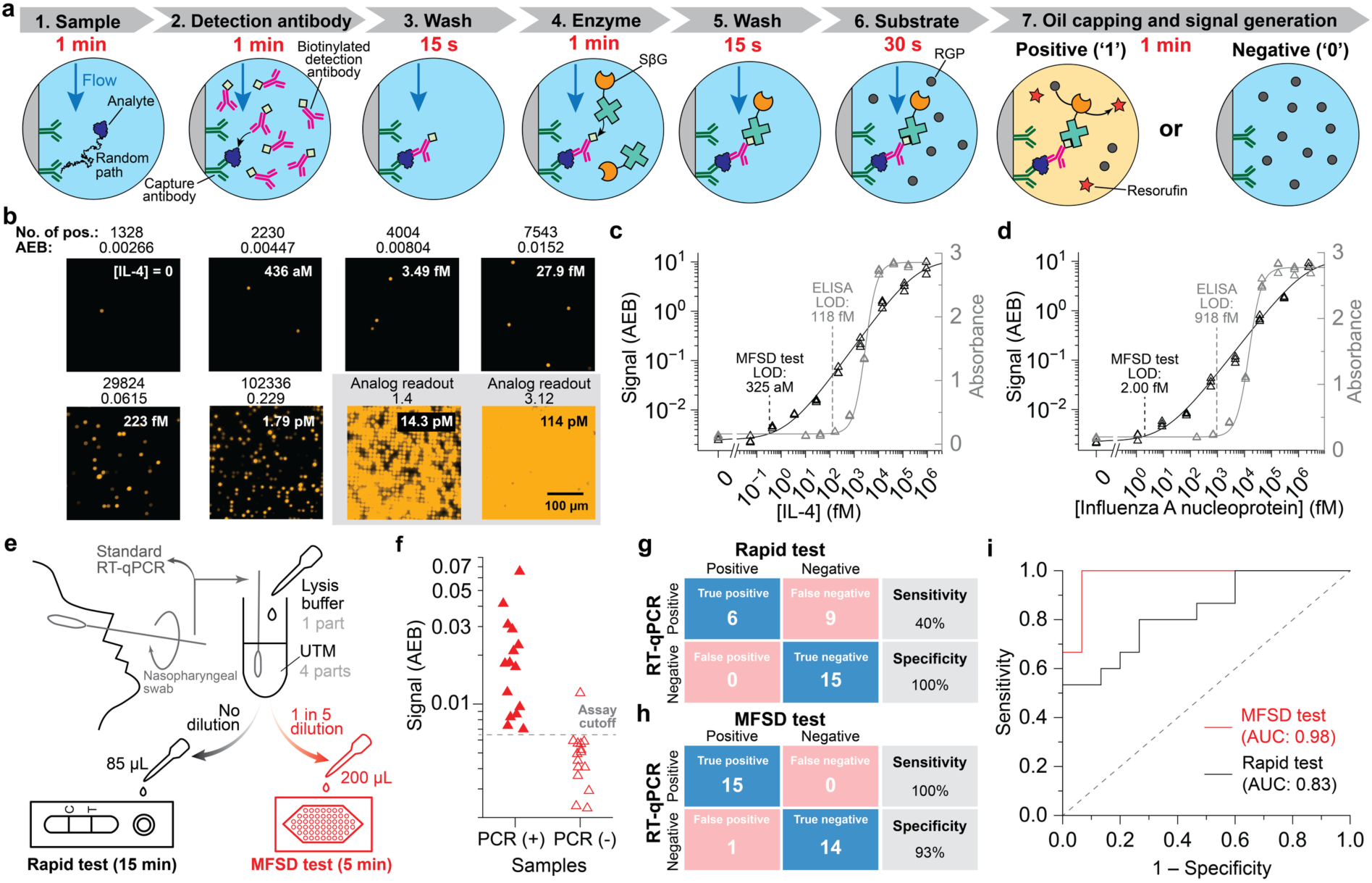
Sandwich MFSD for protein detection and its application to influenza A detection from nasopharyngeal swab samples. **a**, 5-min MFSD test based on a sandwich immunoassay workflow. The sample with the analyte, biotinylated detection antibody, enzyme conjugate, substrate and wash buffers are flowed sequentially for the indicated duration, followed by oil capping of the BATs, signal generation and quantification. **b**, Fluorescence images of the MFSD test against IL-4 with concentrations, positive counts and quantification by calculating the average number of enzymes per BAT (AEB). Quantification is achieved by digital counting (MFSD-D) from ultralow to low concentrations, and by the average signal of the MFSD (MFSD-A) for higher concentrations (greyed area), thereby extending the dynamic range of the test. **c**, Standard curves of 5-min MFSD test for IL-4 and **d**, for influenza A virus nucleoprotein. The standard curves from MFSD test and the benchmark microplate ELISA are shown in black and grey, respectively. The coefficients of variation (CVs) for IL-4 and influenza A nucleoprotein are 14 and 12 %, respectively. All LODs were defined as the concentrations yielding an AEB three standard deviations above the mean AEB of the zero-concentration sample. **e,** Influenza A detection from nasopharyngeal swab samples. Nasopharyngeal swab samples in universal transport medium (UTM) were lysed to release influenza A virus nucleoprotein. The samples were then tested by both a 15-min rapid test without dilution and the 5-min MFSD test after a 5-fold dilution. **f**, MFSD test results of 15 positive and 15 negative samples. **g**, Confusion matrices for the rapid test and **h**, for the MFSD test. **i**, Receiver operating characteristic (ROC) curves for the rapid test and MFSD test. Areas under the curve (AUCs) for the rapid test and MFSD test were evaluated as 0.83 and 0.98, respectively.

To evaluate the potential of the MFSD for clinical translation, we tested 15 influenza A-positive and 15 negative samples obtained via nasopharyngeal swabbing and diagnosed using RT-qPCR (See Extended Data Table 2 for demographics), and compared it to a commercial rapid test. Influenza A nucleoprotein was released by adding lysis buffer and waiting for 5 min, and measured using both the 15-min rapid test and the 5-min MFSD test (Fig. 4e). The rapid test detected 6 out of 15 positives (Extended Data Fig. 5), yielding 40% sensitivity (95% CI, 20–64%), and 100% specificity (95% CI, 80–100%). The MFSD test detected all positives including those missed by the rapid test, with one false positive (Fig. 4f), yielding 100% sensitivity (95% CI, 80–100%) and 93% specificity (95% CI, 70–99%; see Fig. 4g,h for confusion matrices of both tests). Receiver operating characteristic (ROC) curve yielded an area under the curve (AUC) of 0.98 and 0.83 for the MFSD and rapid test (Fig. 4i), respectively. In this small cohort, the MFSD test expectedly outperformed rapid tests while matching PCR sensitivity. Larger cohorts and further investigation will be needed to accurately benchmark MFSD tests. These results illustrate the potential of the MFSD as a rapid, ultrasensitive diagnostic test. The intrinsic sensitivity of the MFSD suggests a potential for improved diagnostic sensitivity compared to PCR when considering that there are ∼530 nucleoproteins for 1 genetic copy per influenza A virus.

### Sandwich MFSD test for nucleic acid detection with sensitivity competitive with PCR

We reasoned that the MFSD test could be used for nucleic acid detection. We selected miR-141 as the target analyte because it is small and compatible with sandwich-based detection needed for SβG binding and enzymatic amplification (Fig. 5a), and compared it to polymerase chain reaction (PCR), which serves as a gold standard. The LOD of the MFSD tests was 7.05 fM for a 200 µL sample, again with a 5 min time-to-result (Fig. 5b) which we found to rival the LOD of reverse-transcription quantitative PCR (RT-qPCR) that was 1.93 fM at the background cycle threshold (Ct) of 35 (Extended Data Fig. 6). However, PCR lasted for 2h and operated from ∼ 1 µL of effective sample volume. Droplet digital PCR (ddPCR) was evaluated but did not outperform the MFSD test; inefficient reverse transcription (RT) and numerous false-positive droplets caused by carryover of the RT primer limited sensitivity, failing detection of miR-141 even at 240 fM (data not shown). The MFSD test illustrates that linear enzymatic signal amplification in a microscopic volume (1 pL) with a high turnover rate (∼10^3^ s^−1^ for βG^36^) can detect molecules in < 1 min, thus outperforming the exponential amplification of PCR by thermocycling. Conventional PCR is further limited to ∼1 µL of effective sample volume and separate upstream preparation (e.g., concentration or extraction) is typically required for larger volumes or complex samples to increase sensitivity. In contrast, MFSD sensitivity can be improved by sieving larger volumes, here with up to 5 mL for 25 min sieving at the same flow rate and an improved LOD of 206 aM (Fig. 5c) and also directly analysed spiked miR-141 in human plasma, with responses in 50% plasma showing good agreement with buffer (Extended Data Fig. 7). The MFSD test thus compares favourably to PCR with its simpler protocol and instrumentation, shorter time-to-result, improved LOD with larger volumes, and compatibility with complex biological matrices.

**Fig. 5.**
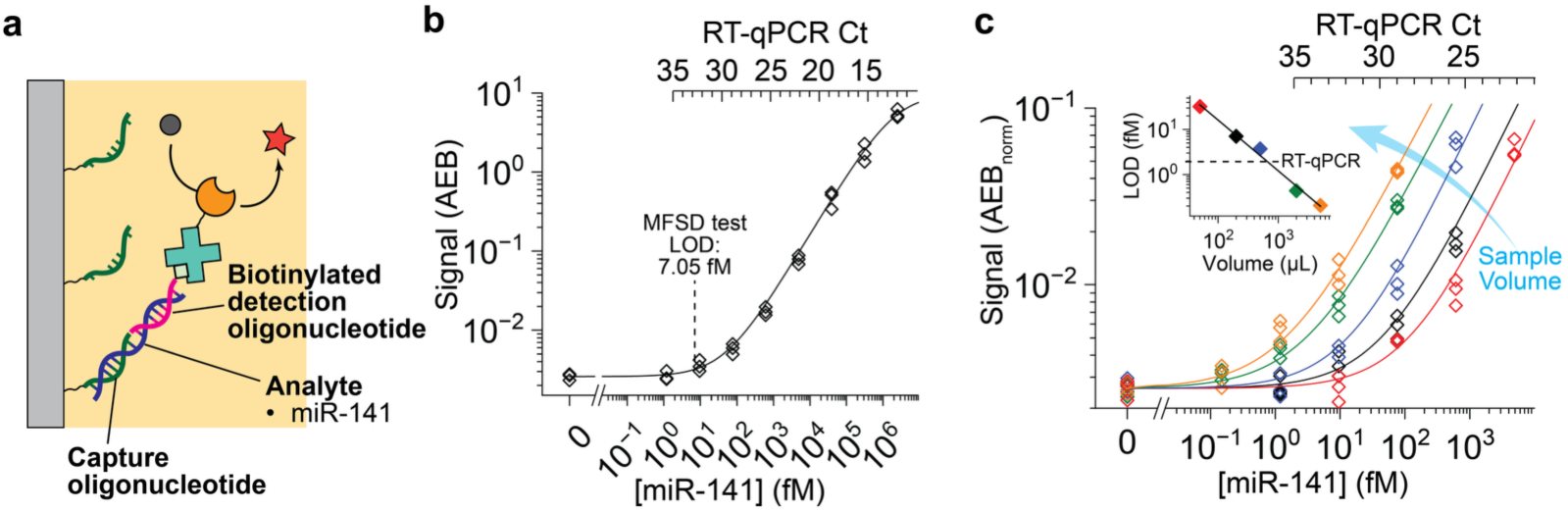
MFSD test detects nucleic acid via sandwich binding much faster than PCR, and is increasingly more sensitive with larger volumes. **a**, Sandwich complex for nucleic acid detections. **b**, Standard curve of 5-min MFSD test for miR-141. The CV is 17%. **c**, Standard curves for volumes of 50 µL (red), 200 µL (black, reference), 500 µL (blue), 2 mL (green) and 5 mL (orange). All samples were flowed at 200 µL min^-1^ followed by the 4 min detection, resulting in total assay times of 4.25, 5, 6.5, 14 and 29 min, respectively. The inset shows LODs for varying sample volumes and corresponding LOD for RT-qPCR as dashed line. All LODs were defined as the concentrations yielding an AEB three standard deviations above the mean AEB of the zero-concentration sample.

## Discussion

The MFSD test uses microfluidic sieving to capture proteins and nucleic acids in 200 µL in one minute with 50% efficiency, and then count individual captured molecules in four more minutes via a ternary sandwich assay and digital detection within partitioned BATs. Digital single molecule assays on microparticles reflect an interpretation of the law of mass action aimed at shifting the binding equilibrium towards more targets being bound by adding many microparticles, and of high capture rates by lengthy incubation according to Langmuir isotherm theory^13,14,22^. The MFSD leads us to reinterpret classical theory via the combination of microscopic confinement and flow, and demonstrates much faster binding than previously conceived, all while capture efficiency *η* is predictable by both ensemble analytical and FEM calculation at Da_I ∼_ 1 and Da_s <_ 1, and confirmed by experiments. However, it is the SBAD that visualizes what a ∼50% capture rate means in a BAT and how it is tuneable by varying flow rates and effective on-rates (*κ*_on)_, as illustrated in the videos. SBAD thus illuminates why 500,000 BATs could sieve a 200-μL-sample in just 1 min.

Conventional macroscopic sieves rely on size-exclusion with pores smaller than the filtrate and are susceptible to clogging and caking, while in molecular sieves pore and molecule size are commensurate^37^. In contrast, in microfluidic sieving introduced here the diameter of the pores (i.e. the BATs) is >1,000 times larger than target analytes, which supports sample flow with hardly any pressure required while simultaneously minimizing the risk of clogging. The MFSD showcases the hitherto untapped potential of vertical flow-through in sensor design that enables rapid capture, and together with partitioning and fast enzymatic signal amplification, make for an overall fast and ultrasensitive test (zeptomolar for a model system), and endow it with a broad dynamic range of seven orders of magnitude and a low CV suitable for quantification. Hence, it improves on the time-to-result of common single molecule immunoassays by well over an order of magnitude, and simultaneously becomes an alternative for nucleic acid detection because it is faster than common nucleic acid tests, and more sensitive with increasing volume, Fig. 6.

**Fig. 6.**
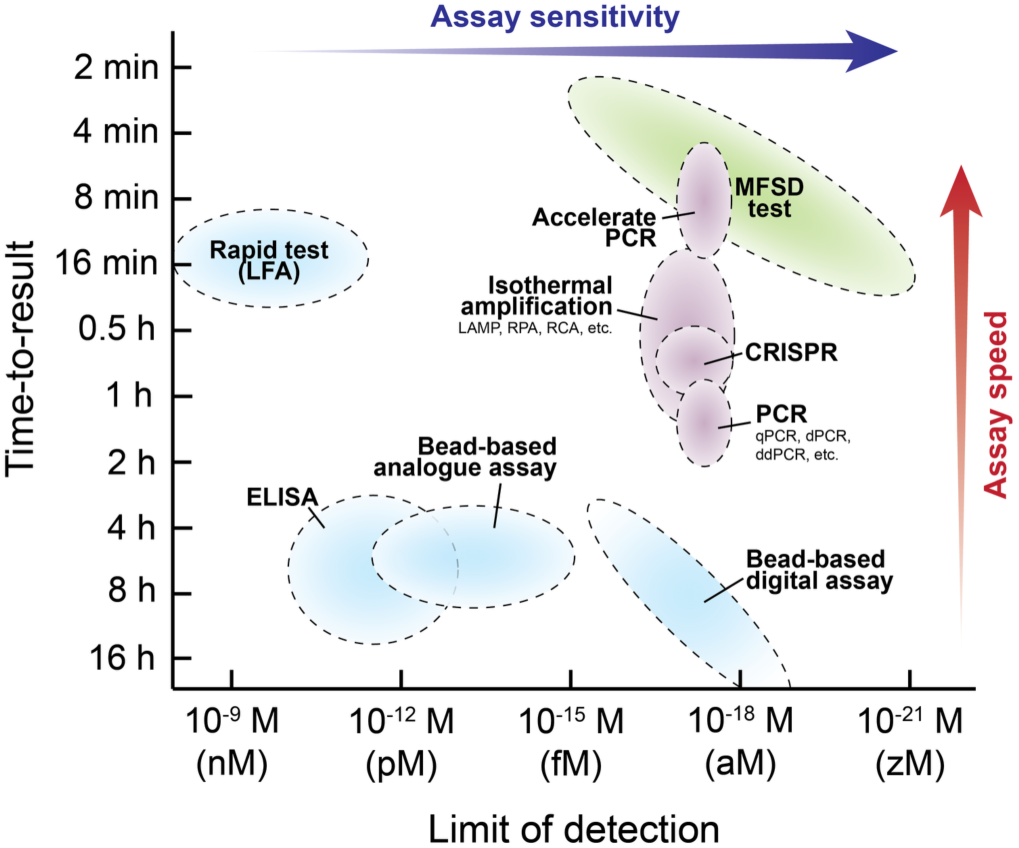
MFSD test benchmarking. The LODs and total assay times of protein assays^4,5,20–24,38,39^ including ELISA, rapid tests, bead-based analogue and digital assays are shown in blue while nucleic acid amplification tests^11^ including PCR, isothermal amplification, and CRISPR-based detection are shown in purple. The current MFSD test is shown as a slanted green oval reflecting its tuneability and trade-off between improving sensitivity further by flowing larger volumes *vs* longer time-to-result.

In this study, we presented the MFSD and its rapid, efficient capture of analytes, and ultrasensitive, *in situ* digital (and analogue) detection in five minutes based on a large array of BATs. The MFSD recasts the trade-off of speed *vs* sensitivity from hours and pM to minutes and aM, all while being simple, highly specific thanks to the sandwich assay configuration and adaptable, thus unlocking amplification-free nucleic acid testing. The MFSD is a thin, isoporous polymer membrane that operates with flow-through and while it was operated manually with various laboratory components here, it is predisposed for integration and for automation, and open to future improvements. The MFSD test, with single molecule sensitivity, short time-to-result, large dynamic and volumetric range and versatility has the potential to redefine the rules of diagnostics and analysis and open multiple new opportunities and applications.

## Materials and methods

### Materials

#### Microfluidic sieve and detector (MFSD) fabrication

Fluorolink MD 700 (Lot #BL2718, Solvay Specialty Polymers USA LLC, Alpharetta, GA, USA), 2-Hydroxy-2-methylpropiophenone (Darocur 1173) (Cat. #405655, lot #MKCD9390, Sigma-Aldrich, Oakville, ON, Canada), Ebecryl® 3708 (Cat. #024709A, lot #NA0275104, Allnex Canada Inc., Ottawa, ON, Canada), Tripropylene glycol diacrylate (TPGDA) (Cat. #027600A, lot #NA0073085, Allnex Canada Inc., Ottawa, ON, Canada), 3,4-Epoxycyclohexylmethyl 3,4-epoxycyclohexanecarboxylate (Cat. #407208, lot #MKCP6704, Sigma-Aldrich, Oakville, ON, Canada), Capa® 3050 (Lot #0000001917, Ingevity UK Ltd, Warrington, Cheshire, UK), Uvacure 1600 (Cat. #028382A, lot #SE103117, Allnex Canada Inc., Ottawa, ON, Canada), Poly(ethylene glycol) diacrylate (PEGDA)-250 (Cat. #475629, lot #MKCS0144, Sigma-Aldrich, Oakville, ON, Canada), diphenyl(2,4,6-trimethylbenzoyl)phosphine oxide (TPO) (Cat. #415952, lot #MKCS4625, Sigma-Aldrich, Oakville, ON, Canada), 2-isopropylthioxanthone (ITX) (Cat. #I067825G, lot #XJHJH-LA, TCI America, Portland, OR, USA), pentaerythritol tetraacrylate (PETTA) (Cat. #408263, lot #MKCR5556, Sigma-Aldrich, Oakville, ON, Canada).

#### MFSD functionalization

N-Acryloxysuccinimide (NAS; Acrylic acid N-Hydroxysuccinimide (NHS) ester) (Cat. #S0814, lot #8QMTA-WX, TCI America, Portland, OR, USA), TPO, Darocur 1173, Dimethyl sulfoxide (DMSO) (Cat. #036480.K2, lot #X11K026, Thermo Fisher Scientific, Waltham, MA, USA), 1H,1H-Heptafluorobutylamine (Cat. #H1300, lot # EMNAH-IR, TCI America, Portland, OR, USA), Biotin-PEG7-amine (Cat. #BP-21622, lot #B103-085, Broadpharm, San Diego, CA, USA), m-PEG2-amine (Cat. #BP-21109, lot #20230322A, Broadpharm, San Diego, CA, USA), Anti-βG (Cat. #Z3783, lot #0000628031, Promega, Madison, WI, USA), Purified anti-streptavidin (Cat. #410501, lot #B294133, Biolegend, San Diego, CA, USA), Capture DNA oligonucleotide for BAT efficiency measurement (5’-/5AmMC12/CAC GCA TCA CCA -3’, Integrated DNA Technologies, Coralville, IA, USA), miR-141 capture oligonucleotide (5’-/5AmMC12/TTT TTT TTC +CAT CT+T TA+C +C -3’, Integrated DNA Technologies, Coralville, IA, USA), Purified anti-human IL-4 (Cat. #500702, lot #B372341, Biolegend, San Diego, CA, USA), Influenza A virus Nucleoprotein antibody (Cat. #GTX41278, lot #822305152, GeneTex, Irvine, CA, USA).

#### Streptavidin-β-galactosidase (SβG) conjugation

Streptavidin protein (Cat. #21122, lot #ZA379579, Thermo Fisher Scientific, Waltham, MA, USA), β-Galactosidase from *Escherichia coli* (Cat. #G5635, lot #0000286885, Sigma-Aldrich, Oakville, ON, Canada), TCO-PEG4-NHS ester (Cat. #BP-22418, lot #B154-066, Broadpharm, San Diego, CA, USA), Methyltetrazine-PEG4-NHS ester (Cat. #BP-22945, lot #B143-046, Broadpharm, San Diego, CA, USA), Glycerol (Cat. #BP229-1, lot #151821, Thermo Fisher Scientific, Waltham, MA, USA).

#### MFSD test for SβG, IL-4, Influenza A nucleoprotein and miR-141

Biotinylated DNA oligonucleotide analyte (5’-/5BiotinTEG/TGG TGA TGC GTG -3’, Integrated DNA Technologies, Coralville, IA, USA), Biotinylated DNA oligonucleotide analyte with double mismatch (5’-/5BiotinTEG/TGG TGT AGC GTG -3’, Integrated DNA Technologies, Coralville, IA, USA), miR-141 (5’-rUrArA rCrArC rUrGrU rCrUrG rGrUrA rArArG rArUrG rG -3’, Integrated DNA Technologies, Coralville, IA, USA), Biotinylated miR-141 detection oligonucleotide (5’-+A+GA +CA+G TG+T TA/3Bio/ -3’, Integrated DNA Technologies, Coralville, IA, USA), Recombinant Human IL-4 Protein (Cat. #204-IL-010/CF, lot #NCL0723062, R&D Systems, Minneapolis, MN, USA), Biotin anti-human IL-4 (Cat. #500804, lot #B360555, Biolegend, San Diego, CA, USA), Influenza A H1N1 (A/Puerto Rico/8/34/Mount Sinai) Nucleoprotein (Cat. #11675-V08B, lot #LC16MY3024, Sino Biological, Wayne, PA, USA), Anti-H7N9 Nucleoprotein (Cat. #11675-R705, lot #HA12DE0602, Sino Biological, Wayne, PA, USA), Resorufin β-D-Galactopyranoside (RGP) (Cat. #R1159, lot #2647827, Thermo Fisher Scientific, Waltham, MA, USA), Fluorinert™ FC-40 (Cat. #F9755, lot #MKCF9430, Sigma-Aldrich, Oakville, ON, Canada), Tris base (Cat. #648311, lot #3782098, Sigma-Aldrich, Oakville, ON, Canada), Sodium chloride (NaCl) (Cat. #S271-500, lot #223105, Thermo Fisher Scientific, Waltham, MA, USA), Magnesium chloride (MgCl_2)_ (Cat. #012315.A1, lot #N22J024, Thermo Fisher Scientific, Waltham, MA, USA), PBS 10x (Cat. #70011-044, lot #3048030, Thermo Fisher Scientific, Waltham, MA, USA), Tween-20 (Cat. #P7949, lot #SLBX0835, Sigma-Aldrich, Oakville, ON, Canada), SuperBlock™ Blocking Buffer (Cat. #37516, lot #AA403418, Thermo Fisher Scientific, Waltham, MA, USA), Nuclease-free water (Cat. #W4502, lot #0000344156, Sigma-Aldrich, Oakville, ON, Canada), RNase Inhibitor, Murine (Cat. #M0314L, lot #0010289386, New England BioLabs, Whitby, ON, Canada), Citrated normal human pooled plasma (Cat. #PL080322MU, lot #BC80322PHP, BioChemed Services, Winchester, VA, USA).

#### Benchmarking RT-qPCR, ddPCR and ELISA

miR-141, TaqMan™ MicroRNA Reverse Transcription Kit (Cat. #4366596, lot #3083491, Thermo Fisher Scientific, Waltham, MA, USA), TaqMan™ MicroRNA Assay (Cat. #4427975, lot #P241216, Thermo Fisher Scientific, Waltham, MA, USA), TaqMan™ Universal Master Mix II, with UNG (Cat. #4440042, lot #3006006, Thermo Fisher Scientific, Waltham, MA, USA), QX200™ droplet generation oil (Cat. #1864005, lot #L006641, Bio-Rad, Hercules, CA, USA), ddPCR™ Supermix for Probes (Cat. #186-3026, lot #64673787, Bio-Rad, Hercules, CA, USA), DG8™ Cartridges for QX200™ Droplet Generator (Cat. #1864008, lot #1000125499, Bio-Rad, Hercules, CA, USA), Nunc™ MaxiSorp™ ELISA Plates, Uncoated (Cat. #423501, lot #B447813, Biolegend, San Diego, CA, USA), Purified anti-human IL-4, Biotin anti-human IL-4, Recombinant Human IL-4 Protein, Influenza A virus Nucleoprotein antibody, Anti-H7N9 Nucleoprotein, Influenza A H1N1 (A/Puerto Rico/8/34/Mount Sinai) Nucleoprotein, Pierce™ High Sensitivity Streptavidin-HRP (Cat. #21130, lot #YE375674, Thermo Fisher Scientific, Waltham, MA, USA), 3,3’,5,5’-Tetramethylbenzidine (TMB) (Cat. #T8665, lot #059K5307, Sigma-Aldrich, Oakville, ON, Canada), Sulfuric acid (H_2S_O_4)_ (Cat. #351293-500, lot #115107, Thermo Fisher Scientific, Waltham, MA, USA), Sodium bicarbonate (NaHCO_3)_ (Cat. #S233-500, lot #071014, Thermo Fisher Scientific, Waltham, MA, USA), Sodium carbonate (Na_2C_O_3)_ (Cat. #S7795, lot #SLBT9993, Sigma-Aldrich, Oakville, ON, Canada), PBS 10x, Tween-20, Bovine serum albumin (BSA) (Cat. #001-000-162, lot #172667, Jackson ImmunoResearch, West Grove, PA, USA).

#### Influenza patient sample analysis

Influenza nasopharyngeal swab samples (American Blood Services DBA, Los Osos, CA, USA), Tris Base, NaCl, DTT (Cat. #R0861, lot #3047435, Thermo Fisher Scientific, Waltham, MA, USA), Halt™ Protease Inhibitor Cocktail (Cat. #78430, lot #3213813, Thermo Fisher Scientific, Waltham, MA, USA), RNase A (Cat. #EN0531, lot #3250577, Thermo Fisher Scientific, Waltham, MA, USA), Glycerol, Triton X-100 (Cat. #BP151-500, lot #051126, Thermo Fisher Scientific, Waltham, MA, USA), Influenza AB + COVID-19 Antigen – 3 in 1 Test (Cat. #COF-19CPC25, lot #I2410155, BTNX Inc., Pickering, ON, Canada).

## Methods

### Stochastic Brownian agent dynamics (SBAD) and finite element method (FEM) simulations

Stochastic Brownian agent dynamics (SBAD) operating the flow chart in Fig. S3 was developed on MATLAB v.R2024a (Mathworks, Natick, MA, USA) and its code is available at https://github.com/junckerlab/microfluidic-sieve-and-detector. For a model 100 kDa globular protein analyte, the diffusion coefficient *D* = 80 µm^2^ s^-1^, was estimated by the Stokes-Einstein-Sutherland equation. SBAD was carried out with Monte Carlo methods, for a high number of analytes to obtain the efficiency *η*. Finite element method (FEM) was performed using COMSOL Multiphysics v6.3 (COMSOL, Inc., Burlington, MA, USA) to obtain ensemble solutions of analyte transport and capture in a BAT over a range of *κ*_on a_nd *Ū*. A three-dimensional model reproducing the BAT geometry was constructed, incorporating the laminar flow, transport of diluted species, chemistry, and surface reactions interfaces. The fluid velocity field was obtained by solving the incompressible Navier-Stokes equations with a prescribed mean inlet velocity, *Ū*, and no-slip boundary conditions at the channel walls. Analyte transport was modelled by convection and diffusion, with a diffusion coefficient of *D*. Surface capture was modelled as a reversible binding reaction between the analyte and immobilized capture probes, with the on-rate *k*_on,_ off-rate *k*_off a_nd surface density of capture probe *Γ*. The capture efficiency, *η*, was determined from the average analyte concentrations at the inlet and outlet planes of the BAT. Based on concentration ergodicity, binding probability *P*_ab i_n SBAD was calibrated by benchmarking against the FEM ensemble solutions (See Supplementary Information for details).

### Fabrication and functionalization of MFSDs

4 × 4 cm^2^ membranes with vertical, 8-µm diameter and 20-µm long pores in a rectangular array of a 40 % open ratio were fabricated with MD 700 by moulding as detailed in our previous publication^32^ and the isoporous membrane was used as an MFSD. Scanning electron microscope (SEM) images of the isoporous membrane were collected on a FEI Quanta 450 environmental scanning electron microscope (FE-ESEM). To fabricate MFSDs with 500,000 BATs on a 3D-printed carrier shown in Fig. 1b, the membranes were cut into 1.3 × 1 cm^2^ pieces, and attached to a carriers 3D-printed with PEGDA-based ink (PLInk)^40^ by using uncured PLInk as a glue to bond the membrane piece and the carrier upon UV exposure. To ensure that the top and bottom surfaces of the MFSDs remained free of capture probe and hydrophobic, the MFSDs were functionalized by the float-flip-float method (Extended Data Fig. 3). First, for NHS-activation, the MFSDs were floated top side-up on 100 mM acrylic acid NHS ester with 10 mM TPO and 2 % (v/v) Darocur 1173 in DMSO, and exposed to UV using a UV Chamber (IntelliRay 600, Uvitron International, Inc., West Springfield, MA, USA) at 70 % power (∼ 120 mW cm^-2^) for 5 min. This floating setup prevented the top surface of the MFSDs from being activated with NHS. The MFSDs were then washed once with DMSO, once with 2-propanol, and then three times with PBS under sonication. The NHS-activated MFSDs were floated top side-down on a solution containing a capture probe (*i.e.,* biotin, antibody and nucleic acid) for 2 h, immobilizing the capture probes only inside the BATs as the top is not NHS-activated and the bottom is not in contact with the solutions. After washing the MFSDs three times with PBS for 1 min each, the MFSDs were submerged into 50 mM 1H,1H-heptafluorobutylamine in PBS for 30 min, quenching unreacted NHS. After 3 washes for 1 min each, the functionalized MFSDs were blocked with SuperBlock™ Blocking Buffer for 1 h before use.

For functionalization with biotin, MFSDs were incubated in a 1 mM biotin in PBS. To vary the surface coverage of biotin on the BATs, mPEG2-amine was mixed with biotin-PEG7-amine in varying ratios while maintaining a total molarity of 1 mM for the mixture. For capture nucleic acid immobilization, amine-modified capture DNA oligonucleotides were used at concentrations of 1 µM for BAT capture efficiency measurements and at 2 µM for miR-141 detection, both prepared in PBS 10x. For antibody functionalization, MFSDs were incubated with a 10 µg ml^-1^ antibody solution prepared in PBS. For quality control, one of the functionalized MFSDs in a batch was fluorescently labelled and inspected under a microscope (Fig. S5b,d,f). Fluorescent labelling was performed using Streptavidin, Alexa Fluor™ 647 Conjugate (Cat. #S32357, lot # 2836838, Thermo Fisher Scientific, Waltham, MA, USA) for biotin-coated surface and nucleic acid-coated surface via the biotinylated analytes or the detection probe, and Goat anti-Mouse IgG (H+L) Cross-Adsorbed Secondary Antibody, Alexa Fluor™ 647 (Cat. #A21235, lot #2482945, Thermo Fisher Scientific, Waltham, MA, USA) for antibody-coated surfaces.

### SβG conjugation

Streptavidin and βG were conjugated by click chemistry^41^. The βG solution was purified by anion exchange chromatography (proFIRE, Dynamic Biosensors GmbH, Munich, Germany) to remove protein aggregates^36^. 10 mg ml^-1^ streptavidin and 3 mg ml^-1^ purified βG solutions were then reacted with TCO-PEG4-NHS ester at 0.9 molar equivalents, and methyltetrazine-PEG4-NHS ester at a 15 molar equivalents, respectively, in PBS for 30 min at room temperature. Following a subsequent buffer exchange to PBS using a Zeba™ spin desalting column (Cat. #89882, lot #YK373207, Thermo Fisher Scientific, Waltham, MA, USA), the activated streptavidin and βG were incubated together overnight at 4 °C to form SβG conjugates by click chemistry. The solution was then filtered using Amicon® Ultra Centrifugal Filter, 100 kDa MWCO (Cat. #UFC5100, lot #0000280113, Sigma-Aldrich, Oakville, ON, Canada) to remove free streptavidin. The purified SβG solution in PBS was mixed with glycerol to a final concentration of 50% (v/v), then supplemented with 5 mM MgCl_2 a_nd stored at –20 °C. SβG concentration was evaluated by absorbance at 280 nm measured with a NanoDrop™ 1000 Spectrophotometer (Thermo Scientific, Waltham, MA USA). The molecular ratio of streptavidin to βG in the conjugated SβG was analysed by performing SDS-PAGE and native PAGE using Mini-PROTEAN® TGX™ precast gels (Cat. #4561096DC, lot #64612653, Bio-Rad, Hercules, CA, USA). It was found that on average, the SβG conjugate contained one βG unit conjugated to 1.23 streptavidin molecules, with an average molecular weight of 530 kDa. Based on Poisson statistics, 29.2% molecules lack streptavidin and consist solely of βG, which was accounted for in the analysis, while free streptavidin had been removed previously via filtration.

### Biotinylation of Influenza A nucleoprotein detection antibody

The detection antibody for Influenza A nucleoprotein (Anti-H7N9 Nucleoprotein) was biotinylated in-house using the EZ-Link™ NHS-PEG4 biotinylation kit (Cat. #21455, lot # YF376472, Thermo Fisher Scientific, Waltham, MA, USA). NHS-PEG4-biotin was prepared at 10 mM in water and added to the antibody solution at a 50-fold molar excess. After 30 min incubation at room temperature, the mixture was buffer-exchanged into PBS using a Zeba™ spin desalting column. The degree of biotinylation was quantified using the standard HABA (4’-hydroxyazobenzene-2-carboxylic acid)/avidin assay. The biotinylated antibody solution was mixed 1:10 into the standard HABA/avidin assay solution containing 0.5 mg/mL avidin (Cat. #1862423, lot #XJ364134, Thermo Fisher Scientific, Waltham, MA, USA) and 300 µM HABA (Cat. #1854180, lot #YB364762, Thermo Fisher Scientific, Waltham, MA, USA) in PBS, and absorbance at 500 nm was then measured using NanoDrop™ 1000 spectrophotometer for quantification of biotin.

### Measurement of capture probe surface density and kinetic constants

The surface density of biotin was measured using the HABA/avidin assay. The pristine and biotin-functionalized MFSDs were each placed in 10 µL of the standard HABA/avidin assay solution, and absorbances of the solution at 500 nm were measured using NanoDrop™ 1000 spectrophotometer to evaluate the depletion of the avidin-HABA complex caused by the surface-immobilized biotin. Considering that the surface binding of each avidin molecule with the tetravalent binding capacity to HABA results in the loss of four HABA molecules in the solution, the estimated biotin density was corrected by dividing by a factor of four. The surface density of biotin was modulated by ratiometric incorporation of polyethylene glycol (PEG) spacers. To determine the surface densities of antibodies, 200 µL of 10 µg mL^-1^ antibody solutions were incubated with and without an NHS-activated MFSD under the same conditions used for the MFSD functionalization. Then, concentrations of the antibody solutions were quantified using the microBCA protein assay kit (Cat. #23235, lot #YC366116, Thermo Fisher Scientific, Waltham, MA, USA) measuring absorbance at 562 nm with NanoDrop™ 1000 spectrophotometer. The difference in concentration between the two conditions was attributed to antibody immobilization on the BAT surface. The surface density of the capture nucleic acid was measured by depletion of fluorescent reporter DNA oligonucleotide complementary to the capture probe. The pristine and functionalized MFSDs were incubated in 10 µL of a 24 nM complementary fluorescent probe (5’-/56-FAM/TGG TGA TGC GT/36-FAM/ -3’, Integrated DNA Technologies, Coralville, IA, USA) in 10 mM Tris, pH 8.0, 1 M NaCl and 0.05% Tween 20 in nuclease-free water (TNT) for 3 min. Reporter concentrations in solution were measured using NanoDrop™ 3300 Fluorospectrometer (Thermo Scientific, Waltham, MA, USA). The concentration difference between the two conditions was attributed to the surface-immobilized capture nucleic acid.

While kinetic constants of biotin and nucleic acid capture probes were obtained from the literature^42,43^, those of anti-streptavidin and anti-βG antibodies were measured by a standard SPR protocol on Sierra SPR®-24 Pro (Bruker, Billerica, MA, USA) with high-capacity amine sensors (Cat. #1814895, lot #0823.a, Bruker, Billerica, MA, USA). The gold surface coated with carboxymethylated dextran was activated by injecting 400 mM EDC (Cat. #03449, lot #BCCG7233, Sigma-Aldrich, Oakville, ON, Canada) and 50 mM NHS (Cat. #24500, lot #XA342567, Thermo Fisher Scientific, Waltham, MA, USA) in water at 5 µL min^-1^ for 420 s, followed by washing with PBS with 0.05% Tween-20 (PBST) at 5 µL min^-1^ for 60 s. Antibodies (50 µg ml^-1^ in 10 mM acetate, pH 5) were immobilized by injection at 5 µL min^-1^ for 600 s. Unreacted NHS esters were quenched by 1M ethanolamine (pH 8.5; Cat. #E9508, lot #SLBR1870V, Sigma-Aldrich, Oakville, ON, Canada) in water at 10 µL min^-1^ for 420 s, followed by a final PBST wash at 10 µL min^-1^ for 240 s. For kinetic analysis, SβG in PBST was injected at 25 µL min^-1^ for 480 s (association), followed by PBST alone at 25 µL min^-1^ for 1,000 s (dissociation). Sensorgrams were analysed using standard kinetic modelling to extract *k*_on a_nd *k*_off v_alues.

### MFSD test experimental setup

As detailed in Fig. 3a, an MFSD was mounted on a PDMS housing cast against a 3D-printed mould. A programmable multi-channel syringe pump (Base module cat. #NEM-B004-01 B, dosing unit cat. #NEM-B002-02 B, Cetoni GmbH, Korbussen, Germany) connected to the PDMS housings via Tygon E-3603 tubing drove the sequential flow for MFSD test. At the end of each flow step, the liquid was withdrawn until the bulk liquid above the MFSD was cleared. The pump was then briefly stopped (<5 s) while the BATs remained filled with the previous liquid, the next liquid was manually loaded on top of the MFSD, and programmed withdrawal was resumed. The MFSDs were then submerged into a bath filled with neat Fluorinert™ FC-40 for 1-min signal generation from 100 µM RGP in PBS with 1 mM MgCl_2._ The MFSDs were placed on an empty petri dish and imaged on an inverted fluorescence microscope (Ti2, Nikon, Tokyo, Japan) equipped with an sCMOS camera (Prime 95B, Teledyne Photometrics, Tucson, AZ, USA) using a 10× objective. Images were acquired with an exposure time of 200 ms. The images were processed using MATLAB with an in-house code (Available at https://github.com/junckerlab/microfluidic-sieve-and-detector) to identify negative (‘0’) and positive (‘1’) BATs, calculate the fraction of positive to total BATs *f*_pos,_ and convert *f*_pos t_o the average number of enzymes per BAT (AEB). At an AEB < 0.5, considering Poisson statistics, the images were digitally processed and *f*_pos w_as converted to AEB by

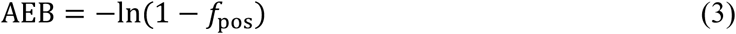

At an AEB > 0.5, the AEB was obtained in an analogue manner as follows.

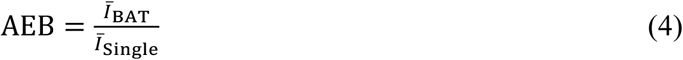

where *Ī*_single i_s an average intensity of single BATs obtained from MFSDs at 0.1 < AEB < 0.5, obtained by averaging intensities globally from multiple batches, and *Ī*_BAT i_s an average intensity of BATs in images under analysis. At least 100,000 BATs (*i.e.*, >20% of the total BATs) were analysed for a statistically robust evaluation of AEBs.

### One-step MFSD test using SβG as the “analyte-detection probe-enzyme”

For one-step MFSD test for BAT efficiency measurement, 200 µL SβG samples in PBST were flowed through MFSDs. Three different concentrations, 2 fM, 400 aM, and 80 aM of SβG, were loaded for biotin-and antibody-functionalized MFSDs. For anti-βG antibody, molarity of the SβG conjugate was adjusted to include βG content, as the anti-βG antibody captures both βG with and without streptavidin. For nucleic acid-functionalized MFSDs, the biotinylated DNA nucleotide analytes, including full complementarity and double mismatch, were pre-coupled with SβG, by spiking SβG at a 200-fold excess (2 nM) into 10 pM analyte. Following 10 min incubation at room temperature, the SβG-oligonucleotide analyte complexes were diluted to final concentrations of 2 fM, 400 aM and 80 aM. After the sample flow, the MFSD was then washed with PBST for 1 min (400 pL at *Ū* = 133 µm s^-1^; *Q* = 6.67 pL s^-1^ per BAT) and filled with 100 µM RGP in PBS with 1 mM MgCl_2 f_or 1 min (400 pL at *Ū* = 133 µm s^-1^; *Q* = 6.67 pL s^-1^ per BAT). Following 1 min signal generation in an FC-40 oil bath, the MFSD was imaged under a microscope and BAT efficiency was obtained as a ratio of the experimental to theoretical AEB. The binding curve for SβG was obtained with 200 µL of SβG standards in SuperBlock™ Blocking Buffer flowed for 1 min (400 pL at *Ū* = 133 µm s^-1^; *Q* = 6.67 pL s^-1^ per BAT) and the same downstream workflow as the BAT efficiency measurement. The experimental limit of detection (LOD) was defined as the concentration yielding an AEB equal to the mean of the blank plus three standard deviations. The Poisson LOD was additionally estimated from three times the Poisson standard deviation of the mean number of BATs classified positive in the blank, which was converted to an input analyte concentration using the measured capture efficiency and 200 µL sample volume.

### Sandwich MFSD test for IL-4 and Influenza A Virus nucleoprotein

For MFSD test for IL-4 and Influenza A nucleoprotein, 200 µL standard samples prepared in SuperBlock™ Blocking Buffer were flowed through MFSDs for 1 min (400 pL at *Ū* = 133 µm s^-1^; *Q* = 6.67 pL s^-1^ per BAT), followed by 10 nM detection antibody in SuperBlock™ Blocking Buffer for 1 min (200 pL at *Ū* = 66.3 µm s^-1^; *Q* = 3.33 pL s^-1^ per BAT). After a wash with PBST for 15 s (400 pL at *Ū* = 531 µm s^-1^; *Q* = 26.67 pL s^-1^ per BAT), 4 pM SβG in SuperBlock™ Blocking Buffer was flowed for 1 min (200 pL at *Ū* = 66.3 µm s^-1^; *Q* = 3.33 pL s^-1^ per BAT). Following a brief wash with PBS for 15 s (400 pL at *Ū* = 531 µm s^-1^; *Q* = 26.67 pL s^-1^ per BAT), the MFSDs were filled with 100 µM RGP in PBS with 1 mM MgCl_2 f_or 30 s (400 pL at *Ū* = 265 µm s^-1^; *Q* = 13.33 pL s^-1^ per BAT). After signal generation in an FC-40 oil bath for 1 min, the MFSDs were imaged under a microscope for AEB readout. All LODs were defined as the concentrations yielding an AEB three standard deviations above the mean AEB of the zero-concentration sample.

### ELISA for IL-4 and Influenza A virus nucleoprotein

Standard ELISA for IL-4 and Influenza A nucleoprotein were performed on Nunc MaxiSorp 96-well plates. Wells were coated with 2 µg mL⁻¹ capture antibody in 50 mM carbonate buffer, pH 9.4 overnight at 4 °C. The wells were washed five times with PBST, each wash 1 min, and blocked with 3% BSA in PBST for 2 h at room temperature. Standard samples were prepared in PBST containing 1% BSA and added to the wells for 2 h incubation at room temperature, followed by five 1 min washes with PBST. Detection antibodies were diluted in PBST with 1% BSA to 2 µg mL⁻¹ for IL-4 and 1 µg mL⁻¹ for Influenza A nucleoprotein and incubated for 1 h at room temperature. The wells were again washed five times with PBST, followed by incubation with streptavidin-HRP diluted 1:50,000 in PBST with 1% BSA for 30 min at room temperature. After a final series of five 1 min washes, TMB substrate was added for signal generation at room temperature and the reaction was stopped using 0.16M sulfuric acid after 10 min. Absorbance was measured at 450 nm using a F129013 Safire plate reader (Tecan, Männedorf, Switzerland). All LODs were defined as the concentrations yielding an AEB three standard deviations above the mean AEB of the zero-concentration sample.

### Sandwich MFSD test for microRNA-141

For MFSD test for miR-141, 200–5000µL miR-141 standard samples prepared in TNT were flowed through MFSDs for 1–25 min (400–10,000 pL at *Ū* = 133 µm s^-1^; *Q* = 6.67 pL s^-1^ per BAT). Then, 10 nM detection oligonucleotide in TNT was flowed through MFSDs for 1 min (200 pL at *Ū* = 66.3 µm s^-1^; *Q* = 3.33 pL s^-1^ per BAT). After a wash with TNT for 15 s (400 pL at *Ū* = 531 µm s^-1^; *Q* = 26.67 pL s^-1^ per BAT), 4 pM SβG in SuperBlock™ Blocking Buffer was flowed for 1 min (200 pL at *Ū* = 66.3 µm s^-1^; *Q* = 3.33 pL s^-1^ per BAT). Following a brief wash with PBS for 15 s (400 pL at *Ū* = 531 µm s^-1^; *Q* = 26.67 pL s^-1^ per BAT), the MFSDs were filled with 100 µM RGP in PBS with 1 mM MgCl_2 f_or 30 s (400 pL at *Ū* = 265 µm s^-1^; *Q* = 13.33 pL s^-1^ per BAT). After signal generation in an FC-40 oil bath for 1 min, the MFSDs were imaged under a microscope for AEB readout. The LOD was defined as the concentrations yielding an AEB three standard deviations above the mean AEB of the zero-concentration sample.

For the buffer-plasma comparison, 2× miR-141 solutions in TNT containing 100 units mL^-1^ murine RNase inhibitor, were mixed with human plasma 1:1 (v/v) immediately before the test, yielding 50% plasma and the final miR-141 concentration matching the buffer standards which also contained 100 units mL^-1^ murine RNase inhibitor. The assay was otherwise performed as described above, except that the final PBS wash for plasma samples was extended from 15 to 45 s (1,200 pL at *Ū* = 531 µm s^-1^; *Q* = 26.67 pL s^-1^ per BAT) to restore hydrophobicity of the top and bottom surfaces of the MFSDs for efficient oil capping. For the buffer-plasma comparison experiments, signal generation in FC-40 was performed for 90 s before imaging. To quantify the effect of the plasma matrix on MFSD response, the relative signal recovery was calculated at each miR-141 concentration using the blank-corrected mean AEB of 50% human plasma relative to that of buffer. Relative signal recovery was calculated as (AEB_plasma –_ AEB_plasma,blank)_ / (AEB_buffer –_ AEB_buffer,blank)_, where AEB_plasma,blank a_nd AEB_buffer,blank a_nd are blank AEBs measured in plasma and buffer, respectively, and AEB_plasma a_nd AEB_buffer a_re AEBs measured at each miR-141 concentration in plasma and buffer, respectively.

### RT-qPCR and RT-ddPCR for microRNA-141

Reverse transcription (RT) was carried out using a stem-loop RT primer to generate elongated cDNA from the 22-nt miR-141. Briefly, 5 µL of miR-141 standard solution was added into a 10 µL RT reaction mix containing the TaqMan™ RT primer. The RT reaction was conducted on Biometra T-Gradient thermocycler (Analytik Jena, Germany) under the thermal cycling conditions as follows: 30 min at 16 °C (annealing), 30 min at 42 °C (reverse transcription), and 5 min at 85 °C (enzyme inactivation) and stored at -20°C (long term storage). Real-time PCR of the miR-141 RT product was performed following the manufacturer’s standard procedure. 0.67 µL of the RT product was added to 9.34 µL of the PCR reaction mix containing PCR primers and TaqMan™ probe. Reactions were initiated with enzyme activation at 95 °C for 10 min, followed by 40 amplification cycles of 95 °C for 15 s (denaturation) and 60 °C for 60 s (annealing/extension).

Thermal cycling and real-time fluorescence recording were performed using a CFX Connect real time PCR machine (Bio-Rad, Hercules, CA, USA). Ct value from the no template control was used to set the Ct cutoff. For ddPCR, 1.34 µL of the miR-141 RT product was added to 18.66 µL of ddPCR mix and loaded into the droplet generator cartridge of a QX200 Droplet Generator (BioRad, USA) along with 65 µL of ddPCR droplet generation oil to generate 37.5 µL of droplets. PCR reaction was initiated at 95 °C for 10 min, followed by 35 amplification cycles of 95 °C for 15 s and 60 °C for 60 s using C1000 Touch Thermal Cycler (Bio-Rad, Hercules, CA, USA). Signal readout was carried out on a QX200 Droplet Reader (Bio-Rad, Hercules, CA, USA).

### Influenza A nasopharyngeal sample analysis

15 influenza A-positive and 15 negative nasopharyngeal swab samples were collected by American Blood Services DBA under an appropriate institutional review board (IRB) protocol (IRB approval ID: 45469786) and purchased. The lysis of influenza A nucleoprotein was performed by mixing the nasopharyngeal swab samples in universal transport medium (UTM) with lysis buffers at a ratio of 1:1:8 (buffer A:buffer B:sample) and incubated for 5 min at room temperature. Buffer A contained 500 mM Tris, 150 mM NaCl, 10 mM DTT, 20 mM EDTA, 10× protease inhibitor cocktail, 1 mg ml^-1^ RNase A, and 50% (v/v) glycerol in water and buffer B consisted of 10% (v/v) Triton X-100 and 150 mM NaCl in water. To perform rapid tests, 85 µL of the processed sample was loaded on Influenza AB + COVID-19 Antigen – 3 in 1 Test. At 15 min, the rapid tests were imaged, and the intensities of the test lines relative to the background were quantified using ImageJ software. To perform MFSD test, the processed sample was diluted 1:5 in SuperBlock™ Blocking Buffer, and 200 µL of the diluted sample was analysed based on the detection method described above.

## Extended Data

**Extended Data Fig. 1.**
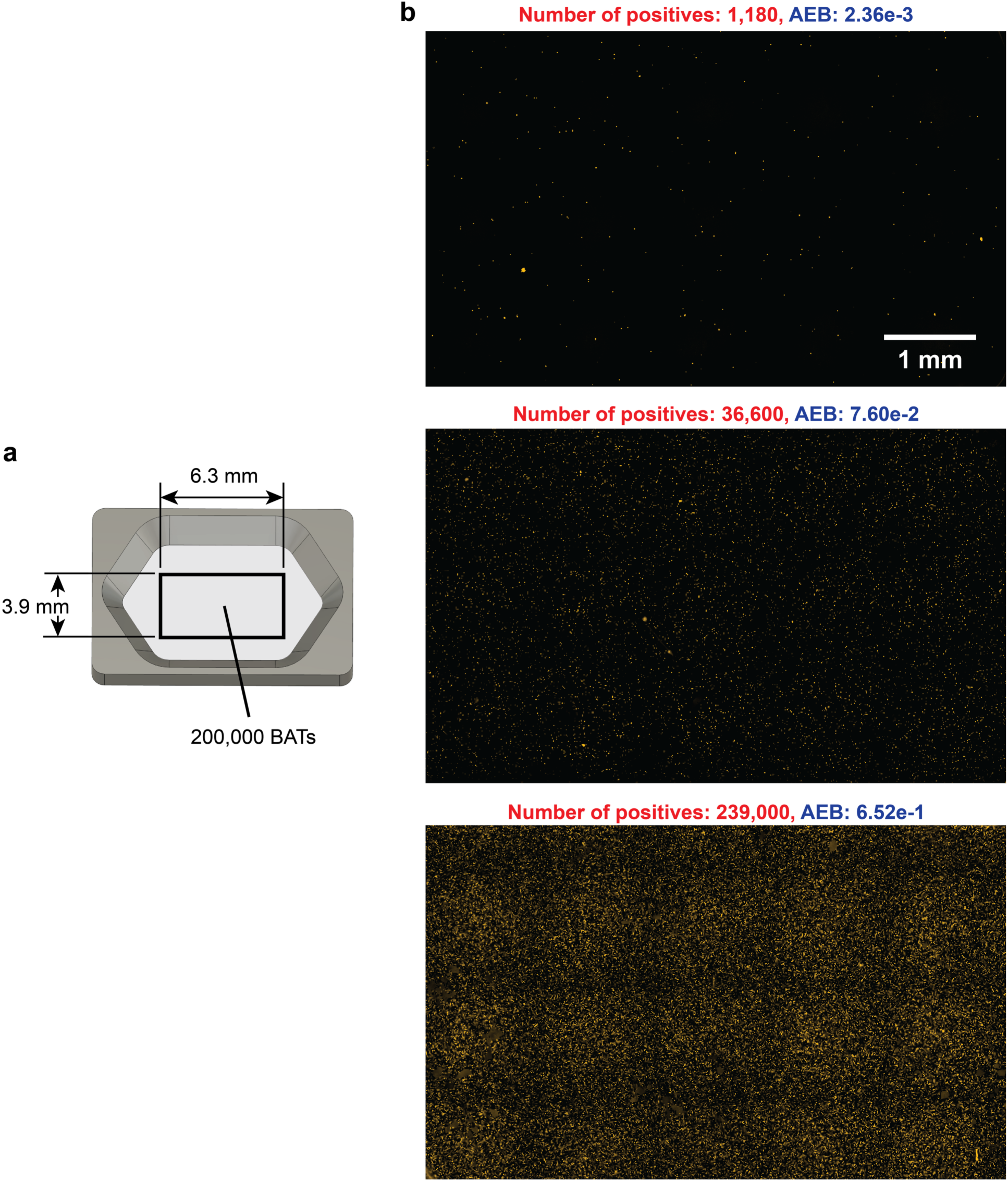
Large-area fluorescent images of MFSDs with detected analytes. **a**, MFSD with an outline of a 6.3 × 3.9 mm^2^ area encompassing 200,000 BATs of the total 500,000 BATs. **b**, Images of the area in **a** generated by stitching multiple fields of view. Exemplar images and digital signals for MFSDs with different total numbers of positive BATs and the corresponding, calculated AEB.

**Extended Data Fig. 2.**
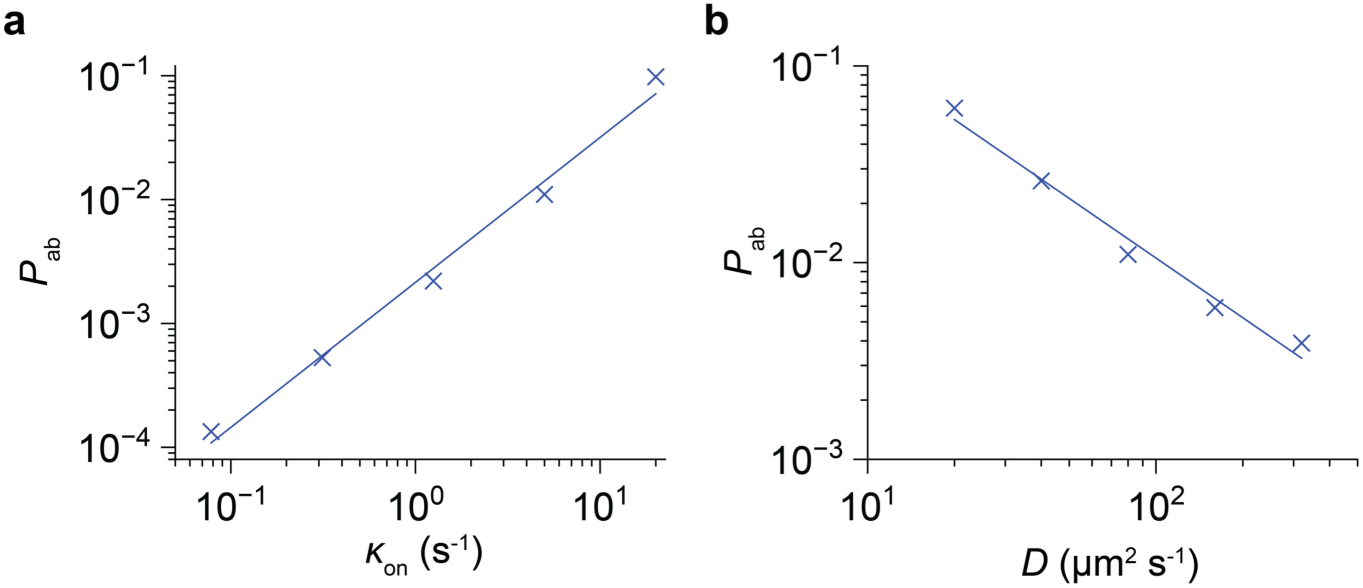
Calibrated affinity binding probability *P*ab. The affinity binding probability, *P*ab was calibrated by sweeping *P*ab until the Monte Carlo estimates of *η* matched the deterministic FEM benchmarks within a 95% confidence interval. Calibration was performed at a fixed flow rate *Ū* = 10 µm s^-1^. **a**, *P*ab calibrated for varying *κ*on at *D* = 80 µm^2^ s^-1^ which is a 100 kDa model globular protein analyte shows a linear relationship (R^2^ = 0.99). **b**, *P*ab calibrated for *D* at *κ*on = 5 s^-1^ shows linear relationship with *κ*on (R^2^ = 0.98).

**Extended Data Fig. 3.**
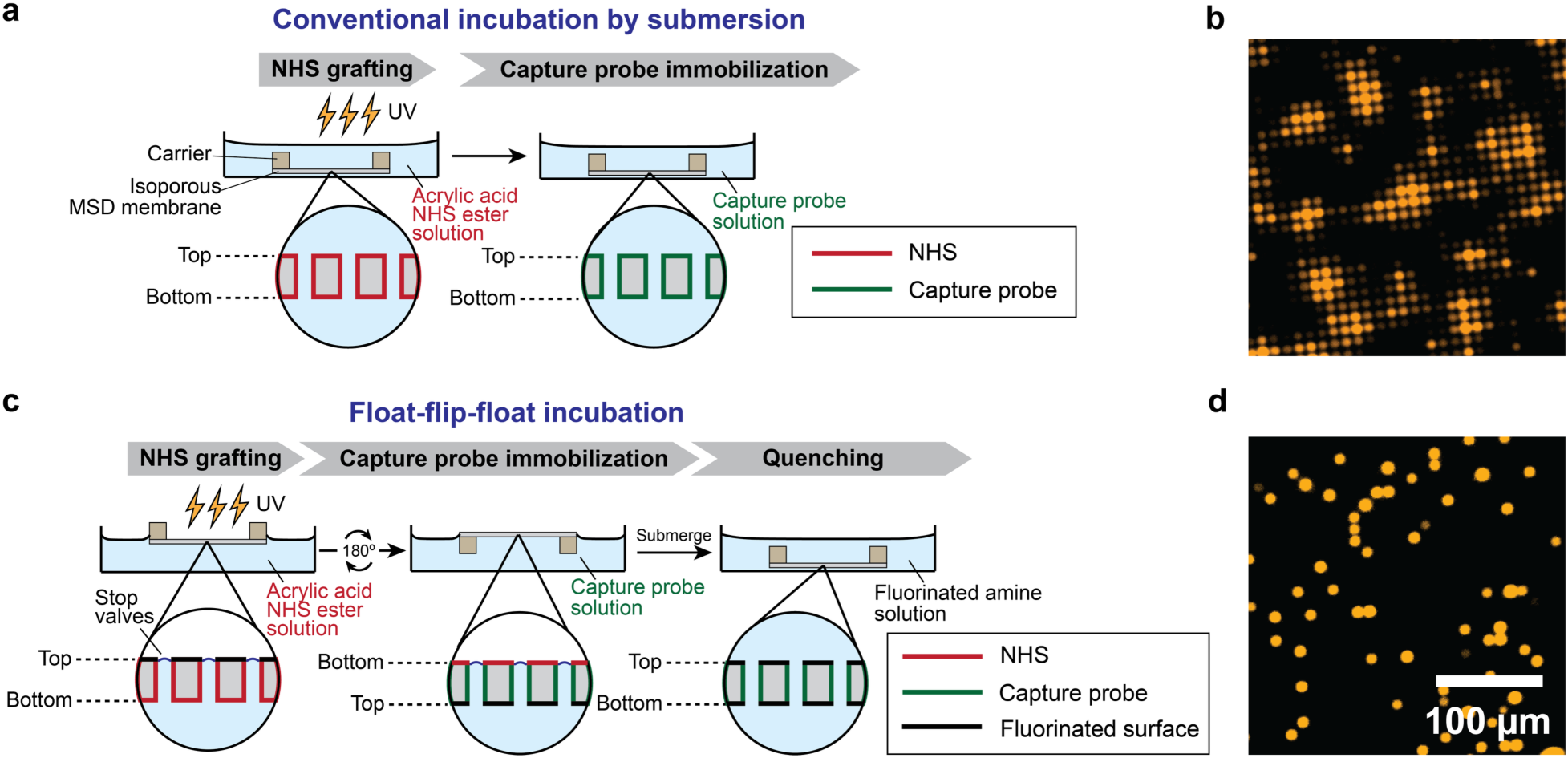
MFSD functionalization by submersion and by float-flip-float. **a**, Conventional incubation of MFSD membranes by submersion in the functionalization solutions, results in capture probes immobilized not only in the pores (i.e. BATs) but also on the top and bottom surfaces of the MFSD, leading to general hydrophilicity. **b**, Ineffective partitioning of BATs due to the hydrophilic top and bottom surfaces causes inter-BAT fluorophore leakage, compromising digital readout. **c**, Float-flip-float incubation. MFSDs are floated by surface tension with BATs (pores) each acting as a stop valve so that NHS-ester groups are only grafted to the ‘bottom surface’ and inside the BATs upon UV light exposure; the ‘top surface’ remains pristine. The membrane is washed, flipped, and floated on a solution containing capture probe molecules with amine groups. Again, the solution contacts the ‘top surface’ (now positioned at the bottom) lacking NHS ester and the inside of the BATs grafted with NHS-ester, resulting in capture probe immobilization exclusively in the BATs. After washing, residual NHS-ester on the ‘bottom surface’ are quenched with 1H,1H-heptafluorobutylamine (*i.e.* fluorocarbon-amine), thus restoring double-sided hydrophobicity. **d**, MFSD assay shows digital signals confined to single BATs, confirming the effective partitioning of the BATs.

**Extended Data Fig. 4.**
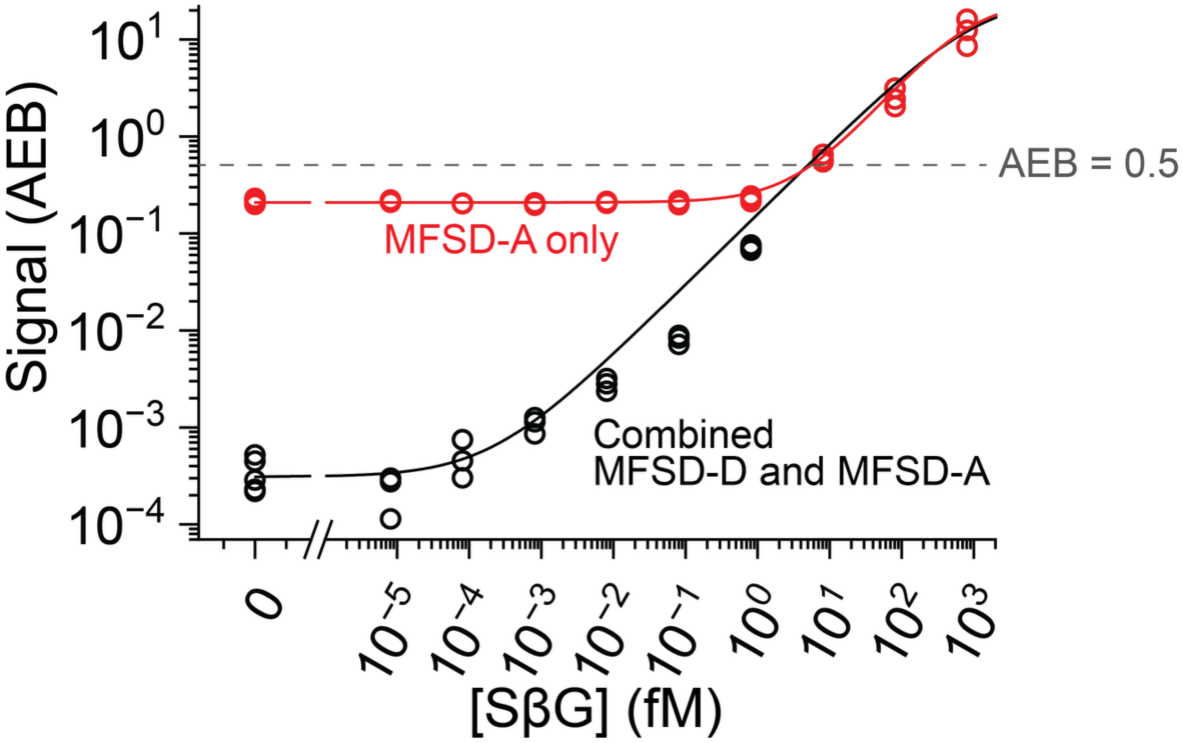
Comparison of “combined MFSD-D and MFSD-A” and “MFSD-A only” readouts. The same MFSDs were read out by the “combined MFSD-D and MFSD-A” and the “MFSD-A only” readouts. The response read out by the MFSD-A only begins to deviate from the combined readout at AEB ≲ 0.5 and approaches its background at an AEB of ∼2.2 × 10^-1^, whereas digital counting unlocks quantification to lower SβG concentrations at the background AEB of 3.4 × 10^-4^ AEB. Thanks to the smooth transition between digital and analogue readouts at AEB = 0.5, the “combined MFSD-D and MFSD-A readout” enables extension of the MFSD dynamic range.

**Extended Data Fig. 5.**
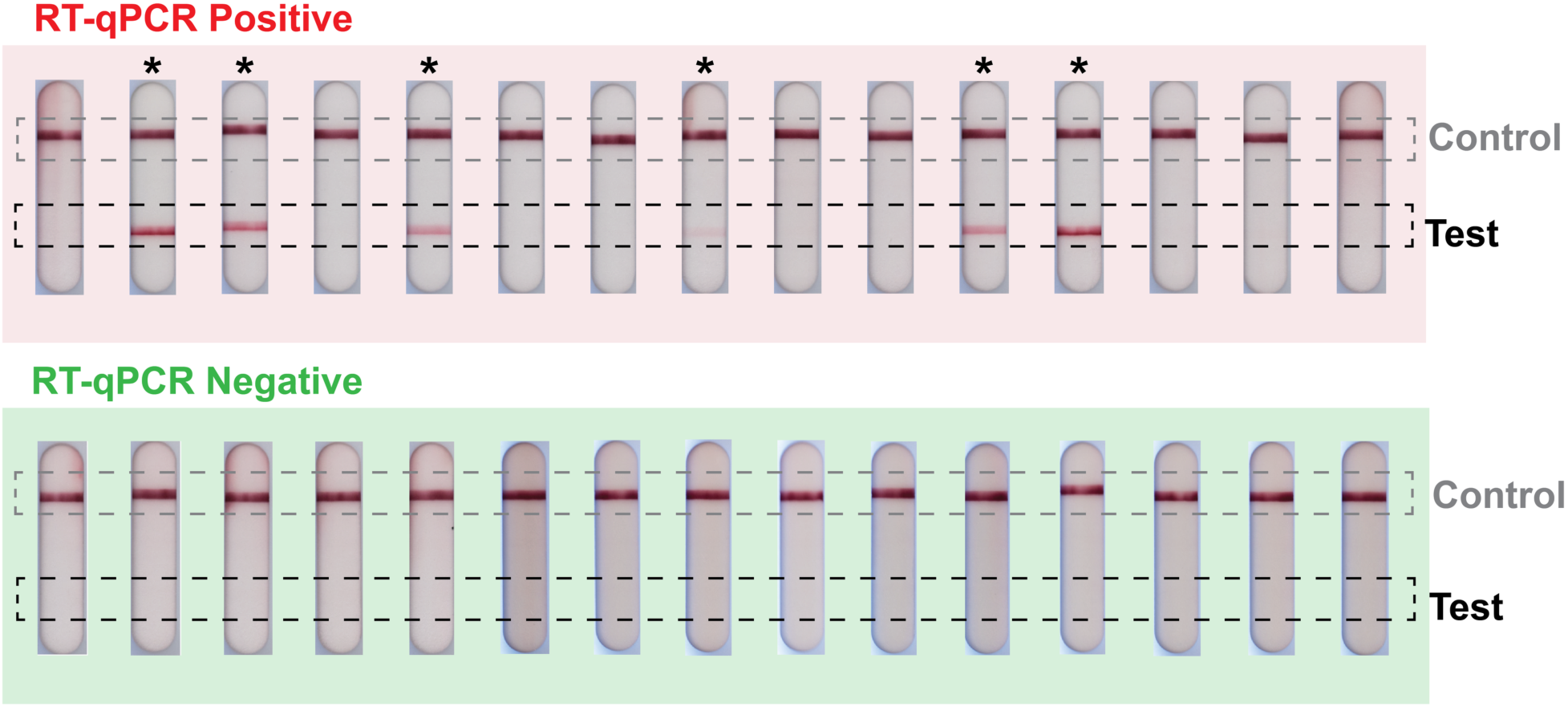
Influenza A rapid tests. Nasopharyngeal swab samples of 15 positive (red panel) and 15 negative samples (green panel) as determined by RT-qPCR were tested using an influenza A rapid test. The rapid test was run for 15 min and then imaged. A visible test line indicates a positive result. The rapid test detected 6 of 15 positive samples and all 15 negative samples were found to be negative. ‘*’ indicates positive rapid test results.

**Extended Data Fig. 6.**
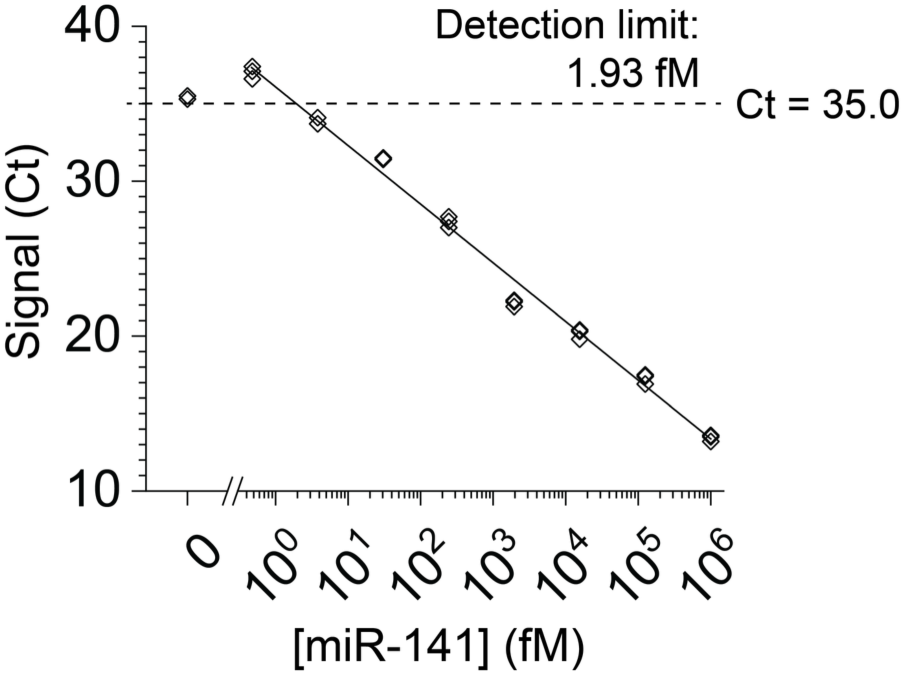
RT-qPCR benchmark for miR-141. RT-qPCR assays for miR-141 were run according to the protocol provided by the vendor, using serially diluted standard samples and no-template control as a negative control. The cut-off Ct was defined as the Ct three standard deviations below the mean assay background.

**Extended Data Fig. 7.**
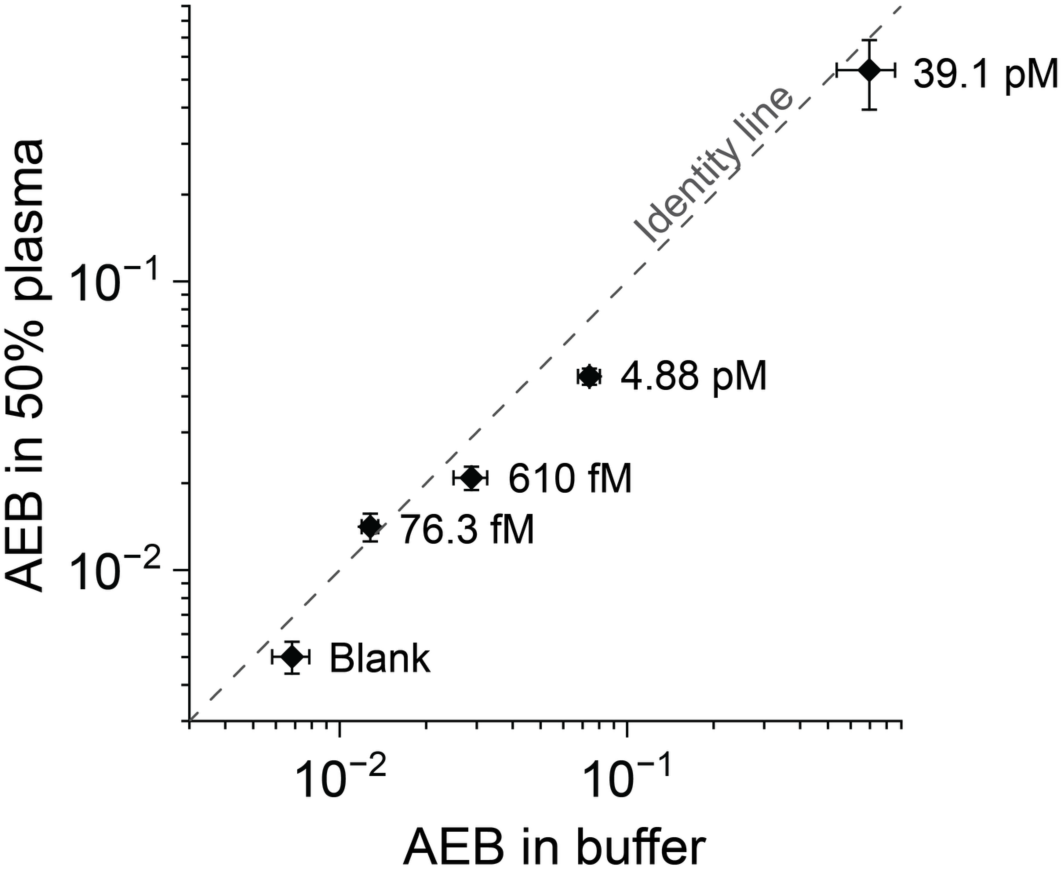
MFSD responses to miR-141 in buffer and 50% plasma. Spiked miR-141 in buffer and 50% human plasma in five matched concentrations were measured by the MFSD. Points and error bars show average responses and standard deviations, respectively (*n* = 3). The dashed line represents identity (AEB in 50% plasma = AEB in buffer). Log-log orthogonal distance regression, accounting for standard deviations of both plasma and buffer measurements, yielded a slope of 0.887 with *R*^2^ = 0.987 (Fitted line not shown). Blank-corrected relative signal recoveries in plasma were 153%, 72%, 62% and 78% at 76.3 fM, 610 fM, 4.88 pM and 39.1 pM, respectively, corresponding to a mean relative signal recovery of 91% across the tested concentrations.

**Extended Data Table 1.** On-rates and surface densities of capture probes in the one-step MFSD test. On-rates *k*on of antibodies were measured by SPR (Fig. S4) while those of biotin and nucleic acid were obtained from the literature^42,43^. The surface densities *Γ* of capture probes were determined by measuring either reporter depletion or free capture probe depletion by surface immobilization (Fig. S5). The biotin surface density was modulated by ratiometric incorporation of a spacer, wherein biotin was introduced as biotin-PEG-amine and the spacer as methyl-PEG-amine at NHS chemistry-based surface coating. Reported values are means.

|  | **Biotin** | **Antibody** |  | **Nucleic acid** |  |
|---|---|---|---|---|---|
|  |  | Anti-strep | Anti-βG | No defects | Double mismatch |
| ***k*<sub>on</sub>**<br>**(× 10<sup>3</sup> m<sup>3</sup> mol<sup>-1</sup> s<sup>-1</sup>)** | 3 | 0.455 | 0.365 | 26.6 | 5.30 |
| ***Γ***<br>**(× 10<sup>-8</sup> mol m<sup>-2</sup>)** | 0.00496–4.96 | 2.08 | 1.25 | 0.0365 | 0.0365 |
| ***κ*<sub>on</sub>**<br>**(s<sup>-1</sup>)** | 0.0744–74.4 | 4.73 | 2.28 | 4.85 | 0.967 |

**Extended Data Table 2.** Nasopharyngeal swab sample demographic characteristics. The demographic characteristics were provided by the supplier.

|  | Number of patients | % of patients |
|---|---|---|
| **Sex** |  |  |
| Male | 12 | 40 |
| Female | 18 | 60 |
| **Age** |  |  |
| 18–30 | 12 | 40 |
| 31–40 | 12 | 40 |
| 41–50 | 6 | 20 |

## Acknowledgements

We gratefully acknowledge M. Tabrizian and J.V.L. Nguyen for access to their SPR system and assistance, A. Kamen, J. Fulber and X. Xu for access to their ddPCR system and assistance, D. Liu at the Facility for Electron Microscopy Research of McGill University for help in microscope operation and data collection, C. Yansouni and J. Papenburg for discussions and advice, T. Gervais for proof-reading, Tri-iso and Ingevity for providing CAPA 3050 samples and Allnex for providing Ebecryl® 3708 and Tripropylene glycol diacrylate (TPGDA) samples. This research was supported by The Natural Sciences and Engineering Research Council of Canada (NSERC) Discovery Grant RGPIN-2022-05171, Canada Research Chair in Bioengineering CRC-232159, and McGill University MI4 Emergency COVID-19 Research Funding and Pathy Seed Fund Grant. G.K. acknowledges an FRQNT doctoral research scholarship and McGill BME recruitment and excellence awards. H.S. acknowledges a Canada Graduate Scholarship – Doctoral (CGSD) and a McGill BME recruitment award. M.L.S acknowledges a Vanier Canada Graduate Scholarship and a McGill BME recruitment award.

## Author contributions

Conceptualization: G.K., A.N., D.J.

Methodology: G.K., A.N., D.J.

Investigation: G.K., H.S., M.L.S.

Visualization: G.K., H.S., M.L.S.

Funding acquisition: A.N., D.J.

Supervision: D.J.

Writing – original draft: G.K., A.N., D.J.

Writing – review & editing: G.K., H.S., M.L.S., A.N., D.J.

## Competing interests

G.K. and D.J. are inventors on a report of invention submitted to McGill University and a related PCT patent application (PCT/CA2025/051350). G.K. and D.J. are co-founders and equity holders in Ubiqui Dx. All other authors declare no competing interests.

## Ethics statement

Nasopharyngeal swab samples were obtained from American Blood Services DBA, Los Osos, CA, USA. According to the supplier’s certification, all samples were collected under an appropriate institutional review board (IRB) protocol (IRB approval ID: 45469786).

## Data availability

All data supporting the conclusion of this study are presented in the article and Supplementary Information. Source data will be available upon request.

## Code availability

MATLAB codes for MFSD image processing and Stochastic Brownian agent dynamics (SBAD) simulation are available at https://github.com/junckerlab/microfluidic-sieve-and-detector.

## Additional information

Supplementary Information is available for this paper. Correspondence and requests should be addressed to.

## Notes

### Summary of Updates

Figures and main text updated with a new acronym. Data added and updated.

## References

1 Squires, T. M., Messinger, R. J. & Manalis, S. R. Making it stick: convection, reaction and diffusion in surface-based biosensors. Nat. Biotechnol. 26, 417–426 (2008).

2 Cesaro-Tadic, S. et al. High-sensitivity miniaturized immunoassays for tumor necrosis factor α using microfluidic systems. Lab Chip 4, 563–569 (2004).

3 Yafia, M. et al. Microfluidic chain reaction of structurally programmed capillary flow events. Nature 605, 464–469 (2022).

4 Elshal, M. F. & McCoy, J. P. Multiplex bead array assays: Performance evaluation and comparison of sensitivity to ELISA. Methods 38, 317–323 (2006).

5 Dagher, M. et al. nELISA: a high-throughput, high-plex platform enables quantitative profiling of the inflammatory secretome. Nat. Methods 22, 2375–2385 (2025).

6 Mullis, K. B. & Faloona, F. A. in Methods Enzymol. Vol. 155 335–350 (Academic Press, 1987).

7 Hindson, B. J. et al. High-Throughput Droplet Digital PCR System for Absolute Quantitation of DNA Copy Number. Anal. Chem. 83, 8604–8610 (2011).

8 Zhao, Y., Chen, F., Li, Q., Wang, L. & Fan, C. Isothermal Amplification of Nucleic Acids. Chem. Rev. 115, 12491–12545 (2015).

9 Shen, F. et al. Digital Isothermal Quantification of Nucleic Acids via Simultaneous Chemical Initiation of Recombinase Polymerase Amplification Reactions on SlipChip. Anal. Chem. 83, 3533–3540 (2011).

10 Broughton, J. P. et al. CRISPR–Cas12-based detection of SARS-CoV-2. Nat. Biotechnol. 38, 870–874 (2020).

11 Filchakova, O. et al. Review of COVID-19 testing and diagnostic methods. Talanta 244, 123409 (2022).

12 Engvall, E. & Perlmann, P. Enzyme-linked immunosorbent assay (ELISA) quantitative assay of immunoglobulin G. Immunochemistry 8, 871–874 (1971).

13 Wilson, B. D. & Soh, H. T. Re-Evaluating the Conventional Wisdom about Binding Assays. Trends in Biochemical Sciences 45, 639–649 (2020).

14 Chang, L. et al. Single molecule enzyme-linked immunosorbent assays: Theoretical considerations. Journal of Immunological Methods 378, 102–115 (2012).

15 Rondelez, Y. et al. Microfabricated arrays of femtoliter chambers allow single molecule enzymology. Nat. Biotechnol. 23, 361–365 (2005).

16 Ge, S., Liu, W., Schlappi, T. & Ismagilov, R. F. Digital, ultrasensitive, end-point protein measurements with large dynamic range via brownian trapping with drift. J. Am. Chem. Soc. 136, 14662–14665 (2014).

17 Honda, S., Minagawa, Y., Noji, H. & Tabata, K. V. Multidimensional Digital Bioassay Platform Based on an Air-Sealed Femtoliter Reactor Array Device. Anal. Chem. 93, 5494–5502 (2021).

18 Byrnes, S. A. et al. Wash-Free, Digital Immunoassay in Polydisperse Droplets. Anal. Chem. 92, 3535–3543 (2020).

19 Han, B. H., Park, M., Chung, S. & Kang, J. Y. Evaporation-driven digital ELISA with micro-droplet arrays for ultrafast detection of low-abundance proteins. Biosens. Bioelectron. 292, 118076 (2026).

20 Rissin, D. M. et al. Single-molecule enzyme-linked immunosorbent assay detects serum proteins at subfemtomolar concentrations. Nat. Biotechnol. 28, 595–599 (2010).

21 Rissin, D. M. et al. Simultaneous Detection of Single Molecules and Singulated Ensembles of Molecules Enables Immunoassays with Broad Dynamic Range. Anal. Chem. 83, 2279–2285 (2011).

22 Kan, C. W. et al. Digital enzyme-linked immunosorbent assays with sub-attomolar detection limits based on low numbers of capture beads combined with high efficiency bead analysis. Lab Chip 20, 2122–2135 (2020).

23 Yelleswarapu, V., et al. Mobile platform for rapid sub–picogram-per-milliliter, multiplexed, digital droplet detection of proteins. Proc. Natl. Acad. Sci. U.S.A. 116, 4489–4495 (2019).

24 Cohen, L. et al. Single Molecule Protein Detection with Attomolar Sensitivity Using Droplet Digital Enzyme-Linked Immunosorbent Assay. ACS Nano 14, 9491–9501 (2020).

25 Risken, H. in The Fokker-Planck equation: methods of solution and applications (Springer, 1989).

26 Fullstone, G., Wood, J., Holcombe, M. & Battaglia, G. Modelling the Transport of Nanoparticles under Blood Flow using an Agent-based Approach. Sci. Rep. 5, 10649 (2015).

27 Tracy, M., Cerdá, M. & Keyes, K. M. Agent-based modeling in public health: current applications and future directions. Annu. Rev. Public Health 39, 77–94 (2018).

28 Bird, R. B., Stewart, W. & Lightfoot, E. N. Transport Phenomena second edition. (John Wiley & Sons, 2002).

29 Gervais, T. & Jensen, K. F. Mass transport and surface reactions in microfluidic systems. Chem. Eng. Sci. 61, 1102–1121 (2006).

30 Brown, G. M. Heat or mass transfer in a fluid in laminar flow in a circular or flat conduit. AIChE Journal 6, 179–183 (1960).

31 Deen, W. M. Analysis of Transport Phenomena. (OUP USA, 1998).

32 Hernández-Castro, J. A., Li, K., Meunier, A., Juncker, D. & Veres, T. Fabrication of large-area polymer microfilter membranes and their application for particle and cell enrichment. Lab Chip 17, 1960–1969 (2017).

33 Meunier, A. et al. Gravity-based microfiltration reveals unexpected prevalence of circulating tumor cell clusters in ovarian and colorectal cancer. Commun. Med. 5, 33 (2025).

34 Leirs, K. et al. Bioassay Development for Ultrasensitive Detection of Influenza A Nucleoprotein Using Digital ELISA. Anal. Chem. 88, 8450–8458 (2016).

35 Rao, E., Mikol, V., Li, D., Kruip, J. & Davison, M. Antibodies that bind IL-4 and/or IL-13 and their uses. United States patent (2020).

36 Rissin, D. M., Gorris, H. H. & Walt, D. R. Distinct and Long-Lived Activity States of Single Enzyme Molecules. J. Am. Chem. Soc. 130, 5349–5353 (2008).

37 Peng, Y. et al. Metal-organic framework nanosheets as building blocks for molecular sieving membranes. Science 346, 1356–1359 (2014).

38 Liu, Y., Zhan, L., Qin, Z., Sackrison, J. & Bischof, J. C. Ultrasensitive and Highly Specific Lateral Flow Assays for Point-of-Care Diagnosis. ACS Nano 15, 3593–3611 (2021).

39 Wu, C., Dougan, T. J. & Walt, D. R. High-Throughput, High-Multiplex Digital Protein Detection with Attomolar Sensitivity. ACS Nano 16, 1025–1035 (2022).

40 Shafique, H. et al. High-resolution low-cost LCD 3D printing for microfluidics and organ-on-a-chip devices. Lab Chip 24, 2774–2790 (2024).

41 Liu, F., Liang, Y. & Houk, K. N. Bioorthogonal Cycloadditions: Computational Analysis with the Distortion/Interaction Model and Predictions of Reactivities. Acc. Chem. Res. 50, 2297–2308 (2017).

42 Srisa-Art, M., Dyson, E. C., deMello, A. J. & Edel, J. B. Monitoring of Real-Time Streptavidin−Biotin Binding Kinetics Using Droplet Microfluidics. Anal. Chem. 80, 7063–7067 (2008).

43 Todisco, M., Ding, D. & Szostak, J. W. Transient states during the annealing of mismatched and bulged oligonucleotides. Nucleic Acids Res. 52, 2174–2187 (2024).

