## Supplementary Information for "Microfluidic sieve and detector for rapid ultrasensitive assays with single-molecule sensitivity"

### Supplementary text

#### Supplementary Discussion S1. Analysis of analyte capture in a BAT using governing parameters and dimensionless numbers

To guide the design and simplify analytical calculations, it is important to understand the parameters that govern analyte capture within a micropore forming a BAT. In this section, taking a BAT with a radius $R_{\text{BAT}}$ = 4 μm (*i.e.* a diameter $d_{\text{BAT}}$ = 8 μm) and a length $L_{\text{BAT}}$ = 20 μm, we analyse analyte capture in a BAT with governing parameters and dimensionless numbers. We consider a model 100 kDa globular protein analyte with a diffusion coefficient, $D$ = 80 μm$^2$ s$^{-1}$ flowing at an average flow speed $\bar{U}$ in the range of $10 < \bar{U} < 4000$ μm s$^{-1}$ at an effective on-rate $\kappa_{\text{on}}$ defined as $\kappa_{\text{on}} = 2k_{\text{on}}\Gamma/R_{\text{BAT}}$ [s$^{-1}$] where $k_{\text{on}}$ is the first-order on-rate [m$^3$ mol$^{-1}$ s$^{-1}$] and $\Gamma$ is the surface density of the capture probe [mol m$^{-2}$], in a range of $0.04 < \kappa_{\text{on}} < 100$ s$^{-1}$.

<u>Mass transport governing wall collision in BATs</u>

We first examine the parameters governing mass transport leading to analyte wall collisions within BATs. Entrance lengths in relation to flow profile development are characterized by the Reynolds number, while the overall analyte mass transport including convection and diffusion is described by the Péclet and Graetz numbers. Together, these dimensionless parameters provide a quantitative framework for assessing analyte transport and wall interactions under varying flow conditions.

Analytes are introduced into the BATs via sample flow, where the velocity profile develops from an initial plug flow to a parabolic profile. As the flow field governs mass transport, identifying the entrance length is essential for accurate modelling. The entrance length is determined by Reynolds number defined as

$$\text{Re} = \frac{\bar{U}d_{\text{BAT}}}{\nu} = \frac{2\bar{U}R_{\text{BAT}}}{\nu} \tag{S1}$$

where $\nu$ is a kinematic viscosity [m$^2$ s$^{-1}$]. With the Reynolds number Re, the hydrodynamic entrance length is found as

$$L_{\text{entrance}} = 0.0575 \cdot \text{Re} \cdot d_{\text{BAT}} \tag{S2}$$

At the flow rate range in this study, $10 < \bar{U} < 4{,}000$ μm s$^{-1}$, Re is in the range of $8.97 \times 10^{-5} < \text{Re} < 3.59 \times 10^{-2}$, resulting in the entrance length in the range of 0.0413 nm < $L_{\text{entrance}}$ < 16.5 nm. Given the length of a BAT, $L_{\mathrm{BAT}}$ = 20 μm, the entrance length is negligibly short. Hence, we assume a parabolic flow profile regardless of the length that the sample has flowed through.

Wall collision in a BAT is governed by mass transport consisting of axial convection and radial diffusion. In microfluidics, Péclet number (Pe) has been commonly used to describe the balance between the convective and diffusive transport, using a single characteristic length, which is diameter, $d_{\mathrm{BAT}}$, to capture the local effect of mass transport. By taking the ratio of the characteristic time scales for convection (~ $d_{\mathrm{BAT}}/\bar{U}$) and diffusion (~ $d_{\mathrm{BAT}}^2/D$), Pe is defined as[1-3]:

$$\mathrm{Pe} = \frac{\bar{U} d_{\mathrm{BAT}}}{D} = \frac{2\bar{U} R_{\mathrm{BAT}}}{D} \tag{S3}$$

However, this simplification ignores the anisotropic nature of mass transport in microchannel-based sensors, where the relevant competing mass transports are radial diffusion through the channel diameter (characteristic time ~ $d_{\mathrm{BAT}}^2/D$), and axial convection through the channel length (characteristic time ~ $L_{\mathrm{BAT}}/\bar{U}$). To more accurately describe this interplay, we employ Graetz number (Gz) which explicitly accounts for both length scales and provides a more comprehensive measure of transport balance[1,2]:

$$\mathrm{Gz} = \frac{\bar{U} d_{\mathrm{BAT}}^2}{L_{\mathrm{BAT}} D} = \frac{4\bar{U} R_{\mathrm{BAT}}^2}{L_{\mathrm{BAT}} D} \tag{S4}$$

Both Pe and Gz greater than 1 indicate convection-dominated transport, while those less than 1 correspond to diffusion-dominated regime. In a BAT, a sample containing the model analyte with $D$ = 80 μm² s⁻¹ flowing at $10 < \bar{U} < 4{,}000$ μm s⁻¹, yields Pe > 1, indicating that the convection generally dominates over diffusion at a local point of the BAT. A complementary description is provided by Gz (Fig. S1a), evaluated over the same velocity range. At $\bar{U} < 25$ μm s⁻¹, Gz is less than 1, which means that analytes have sufficient time to migrate toward and interact with the channel wall. However, at $\bar{U} > 25$ μm s⁻¹, Gz is greater than 1, indicating that axial convection dominates, reducing the efficiency of diffusive transport of analytes to the capture surface.

Dynamics of analyte capture governed by reaction and mass transport

Following mass transport leading to analyte wall collisions, we now assess the kinetics of surface capture. The Damköhler numbers compare the rate of reaction to mass transport, providing a quantitative measure of whether analyte binding is transport- or reaction-limited. Taking residence time of the sample in a BAT as a mass transport term, we get first Dahmköhler number $\mathrm{Da_I}$ defined as[1]:

$$\mathrm{Da_I} = \frac{\kappa_{\mathrm{on}} L_{\mathrm{BAT}}}{\bar{U}} = \kappa_{\mathrm{on}} \tau_{\mathrm{res}} \tag{S5}$$

where $\tau_{\mathrm{res}}$ is the average residence time of the sample. For a BAT as one effective sensor, $\mathrm{Da_I}$ serves as a measure of how fully the analyte capture reaction can progress while the sample flows through a BAT. At $\mathrm{Da_I} \ll 1$, the residence time is not sufficient at the given effective surface on-rate $\kappa_{\mathrm{on}}$ and therefore most analyte molecules leave the BAT reaching the BAT efficiency, $\eta \sim 0$, whereas at $\mathrm{Da_I} \gg 1$, analyte capture in the BAT approaches $\eta \sim 1$.

However, $\mathrm{Da_I}$ does not consider that a BAT sensor is based on surface reactions that must consider mass transport from the bulk to the surface. Thus, we consider the surface reaction and the mass transport to the surface and introduce surface Damköhler number $\mathrm{Da_s}$. According to first-order Langmuir kinetics, the reactive flux $J_{\mathrm{r}}$ [mol m$^{-2}$ s$^{-1}$] is obtained as $J_{\mathrm{r}} = k_{\mathrm{on}} \Gamma c_{\mathrm{w}}$. Based on thin film theory, the mass transport to the surface by convection and diffusion is defined as $J_{\mathrm{m}} = k_{\mathrm{c}}(\bar{c} - c_{\mathrm{w}})$, where $k_{\mathrm{c}}$ is the mass transfer rate [m s$^{-1}$], $J_{\mathrm{m}}$ is the mass transport flux [mol m$^{-2}$ s$^{-1}$], and $c_{\mathrm{w}}$ and $\bar{c}$ are the at-wall and local bulk average concentrations [mol m$^{-3}$], respectively. By equating the fluxes, $J_{\mathrm{r}} = J_{\mathrm{m}}$, we obtain the ratio of at-wall vs bulk concentration,

$$\frac{c_{\mathrm{w}}}{\bar{c}} = \frac{1}{1 + \frac{k_{\mathrm{on}} \Gamma}{k_{\mathrm{c}}}} \tag{S6}$$

We introduce a surface Damköhler number ($\mathrm{Da_s}$) defined as the ratio of surface reaction to total mass transport including both convection and diffusion[3] as function of $k_{\mathrm{on}}$ and $\kappa_{\mathrm{on}}$:

$$\mathrm{Da_s} = \frac{k_{\mathrm{on}} \Gamma}{k_{\mathrm{c}}} = \frac{\kappa_{\mathrm{on}} R_{\mathrm{BAT}}}{2 k_{\mathrm{c}}} \tag{S7}$$

and thus simplify Eq. S6 to:

$$\frac{c_{\mathrm{w}}}{\bar{c}} = \frac{1}{1 + \mathrm{Da_s}} \tag{S8}$$

At $\mathrm{Da_s} \gg 1$, the at-wall concentration $c_{\mathrm{w}}$ approaches 0, indicating that the rate of surface affinity binding at the capture surface vastly exceeds the rate of mass transport (i.e., mass transport-limited), whereas $c_{\mathrm{w}} \sim \bar{c}$ at $\mathrm{Da_s} \ll 1$ implies that mass transport of the analytes to the capture surface operates on a shorter time scale than surface binding (i.e., reaction-limited). Fig. S1b,c show that analyte capture in BATs is generally reaction-limited. For the model analyte with $D$ =

80 µm$^2$ s$^{-1}$, the mass transport rate consistently outpaces the binding rate. Specifically, at a typical capture surface with $\kappa_{on}$ = 5 s$^{-1}$, which is in the same order of magnitude across various types of capture probes (i.e. biotin, antibody and nucleic acid in this study), $Da_s$ remains below 1 across a broad range of flow speed (10 < $\bar{U}$ < 4,000 µm s$^{-1}$; Fig. S1b). Even with increasing $\kappa_{on}$ from 0.04 to 100 s$^{-1}$, at a fixed average flow speed of 133 µm s$^{-1}$ (corresponding to 1 min incubation for 200 µL sample with 500,000 BATs), $Da_s$ < 1 for $\kappa_{on} \leq 20$ s$^{-1}$ (Fig. S1c). In our studies, all capture probe types (*i.e.,* biotin, antibodies, and nucleic acids) had a $\kappa_{on}$ < 20 s$^{-1}$, and hence $Da_s$ < 1. We thus conclude that analyte capture in the BATs in this study was primarily governed by the time scale of intrinsic binding kinetics because mass transport time scales were much shorter.

We note that $k_c$ is used to calculate the Sherwood number, $\mathrm{Sh} = 2k_c R_{BAT}/D$, which represents the ratio of total (convection and diffusion) to diffusive mass transport rate[1,2], or conversely $k_c$ can be expressed as:

$$k_c = \frac{\mathrm{Sh}D}{2R_{\mathrm{BAT}}} \tag{S9}$$

For a tube where a sample flows through (*e.g.*, a BAT), Sh can be obtained as a function of Gz as follows[4]:

$$\mathrm{Sh} = 3.66 + \frac{0.0668\mathrm{Gz}}{1+0.04\mathrm{Gz}^{\frac{2}{3}}} \tag{S10}$$

With these relationships, it is possible to relate Da, Sh and Gz numbers, and identify the governing parameters and time scales depending on the applications.

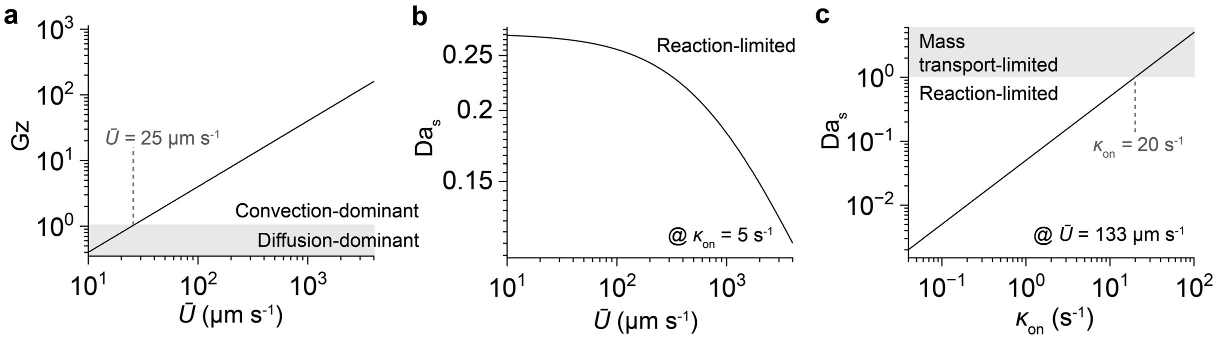


**Fig. S1 | Analysis of analyte mass transport and capture in a BAT with dimensionless numbers.** A model 100 kDa globular protein analyte with a diffusion coefficient $D = 80\ \mu m^2\ s^{-1}$ flows through a BAT with a radius $R_{BAT} = 4\ \mu m$ and a length $L_{BAT} = 20\ \mu m$ with an effective on-rate $\kappa_{on}$ in the range of $0.04 < \kappa_{on} < 100\ s^{-1}$ at a varying flow rate $\bar{U}$ in the range of $10 < \bar{U} < 4{,}000\ \mu m\ s^{-1}$. **a**, Graetz number Gz. Gray shading indicates the region where mass transport is dominated by diffusion. **b**, Surface Damköhler number $Da_s$ with a varying $\bar{U}$ at a fixed $\kappa_{on} = 5\ s^{-1}$. Analyte capture in this condition is reaction limited regardless of $\bar{U}$. **c**, $Da_s$ with a varying $\kappa_{on}$ at a fixed $\bar{U} = 133\ \mu m\ s^{-1}$. Gray shading indicates the region where analyte capture is limited by mass transport.

**Supplementary Discussion S2. Analytical solution of BAT efficiency**

In this section, we derive the analytical solution for BAT efficiency, following the established mathematical formulation and applying it to our specific case[2,5,6]. We use cylindrical coordinates where a point is described by radial distance ($r$) from the origin in the $xy$-plane, the angle ($\theta$) from the $x$-axis in the $xy$-plane and the height ($z$) above or below the $xy$-plane. Throughout the mathematical derivation, we consider an infinitesimal control slice with the thickness of $dz$ (Fig. S2). We consider an ensemble model with the first-order Langmuir kinetics for surface affinity binding, the film theory for mass transport and Reynolds transport theorem for molecular mass balance[7]. We posit that average analyte capture of single molecules in the BATs is equivalent to ensemble models, assuming ergodicity.

Because a BAT is homogeneously coated with a capture probe and thus captures analytes equally throughout all the directions, the concentration field in a BAT is axisymmetric ($\partial/\partial\theta = 0$). For simplicity, we average the local concentration $c$ [mol m$^{-3}$] along $r$ and $\theta$, and get the local bulk average concentration $\bar{c}$ [mol m$^{-3}$], which is a function of $z$ and $t$, $\bar{c}(z, t)$. Because in our experiments, $\Gamma$ is in great excess compared to that of the captured analyte $\Gamma_{\text{captured}}$ [mol m$^{-2}$], the capture surface is not saturated and reaches a constant reaction ($J_r$) and mass-transport ($J_m$) flux ($\partial J_r/\partial t = 0$ and $\partial J_m/\partial t = 0$) and thus reaches a constant-flux steady state with the time-invariant $\bar{c}$ ($\partial\bar{c}/\partial t = 0$).

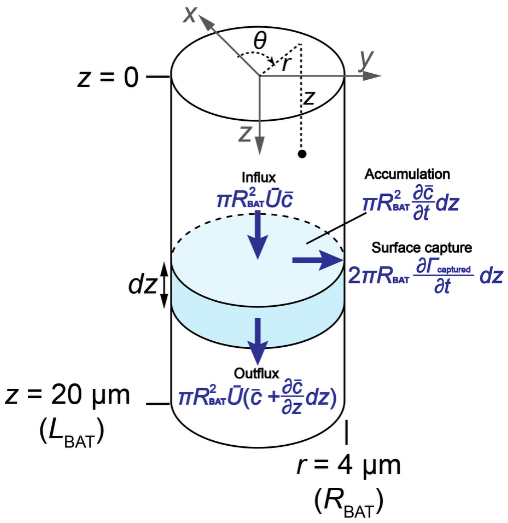


**Fig. S2 | BAT in cylindrical coordinates.** Reynolds transport theorem considers the balance between the molecular accumulation and surface capture along with fluxes that are into and out of the infinitesimal control slice with the thickness $dz$.

The mass transport of the analyte to the wall is considered based on the film theory using a mass-transfer coefficient $k_c$ in Eq. S9. The mass transport flux $J_m$ [mol m$^{-2}$ s$^{-1}$], is driven by the difference between $\bar{c}$ and the at-wall concentration $c_w(z, t)$ [mol m$^{-3}$].

$$J_m = k_c(\bar{c} - c_w) \tag{S11}$$

The affinity binding is considered by first-order Langmuir kinetics which gives reactive flux $J_r$ [mol m$^{-2}$ s$^{-1}$],

$$J_r = \frac{\partial \Gamma_{captured}}{\partial t} = k_{on} c_w (\Gamma - \Gamma_{captured}) - k_{off} \Gamma_{captured} \tag{S12}$$

where $k_{off}$ is an off-rate in [s$^{-1}$]. Due to $\Gamma_{captured} << \Gamma$, Eq. S12 can be re-written as

$$J_r = \frac{\partial \Gamma_{captured}}{\partial t} = k_{on} c_w \Gamma - k_{off} \Gamma_{captured} \tag{S13}$$

At the constant-flux steady state, the concentration field is time-invariant. Hence, the at-wall concentration does not change over time, which means that all the analytes brought to the wall are captured ($J_r = J_m$). Therefore, equating Eqs. S11 and S13, $c_w$ is

$$c_w = \frac{k_c}{k_c + k_{on}\Gamma} \bar{c} + \frac{k_{off}}{k_c + k_{on}\Gamma} \Gamma_{captured} \tag{S14}$$

With substitutions, Eq. S14 is simplified as

$$c_w = \alpha \bar{c} + \beta \Gamma_{captured}, \quad \text{where } \alpha = \frac{k_c}{k_c + k_{on}\Gamma}, \quad \beta = \frac{k_{off}}{k_c + k_{on}\Gamma} \tag{S15}$$

By inserting Eq. S15 into Eq. S13 we find

$$\frac{\partial \Gamma_{captured}}{\partial t} = k_{on} \Gamma \alpha \bar{c} - (k_{off} - k_{on} \Gamma \beta) \Gamma_{captured} \tag{S16}$$

The local bulk average concentration $\bar{c}$, can be mathematically written based on Reynolds transport theorem for the molecular mass conservation, which states the accumulation of the analyte in a control slice balances with influx, outflux and surface capture of the analyte (Fig. S2).

$$\underbrace{\pi R_{BAT}^2 \frac{\partial \bar{c}}{\partial t} dz}_{\text{Accumulation}} = \underbrace{\pi R_{BAT}^2 \bar{U} \bar{c}}_{\text{Influx}} - \underbrace{\pi R_{BAT}^2 \bar{U} \left( \bar{c} + \frac{\partial \bar{c}}{\partial z} dz \right)}_{\text{Outflux}} - \underbrace{2\pi R_{BAT} \frac{\partial \Gamma_{captured}}{\partial t} dz}_{\text{Surface capture}} \tag{S17}$$

As $\partial \bar{c} / \partial t = 0$ in the constant-flux steady state,

$$\bar{U} \frac{\partial \bar{c}}{\partial z} = -\frac{2}{R_{BAT}} \frac{\partial \Gamma_{captured}}{\partial t} \tag{S18}$$

To obtain an analytical solution, we simplify the partial differential equations to ordinary differential equations by taking material derivatives, where we track a control slice travelling through the BAT from the entry at $z = 0$ and the exit at $z = L_{\mathrm{BAT}}$. The total derivative of $\bar{c}$ with respect to $z$ is

$$\frac{d\bar{c}}{dz} = \frac{\partial\bar{c}}{\partial t}\frac{dt}{dz} + \frac{\partial\bar{c}}{\partial z} = \frac{\partial\bar{c}}{\partial z} \quad \text{(S19)}$$

because $\partial\bar{c}/\partial t = 0$, while the total derivative of $\Gamma_{\mathrm{captured}}$ with respect to $z$ is

$$\frac{d\Gamma_{\mathrm{captured}}}{dz} = \frac{\partial\Gamma_{\mathrm{captured}}}{\partial t}\frac{dt}{dz} + \frac{\partial\Gamma_{\mathrm{captured}}}{\partial z} = \frac{1}{\bar{U}}\frac{\partial\Gamma_{\mathrm{captured}}}{\partial t} \quad \text{(S20)}$$

where $dz/dt = \bar{U}$, and $\partial\Gamma_{\mathrm{captured}}/\partial z \approx 0$ because no significant changes in binding occur over the infinitesimal thickness $dz$ of the slice. With Eqs. S19 and S20, partial differential equations, Eqs. S16 and S18 are updated into the following ordinary differential equation:

$$\frac{d\Gamma_{\mathrm{captured}}}{dz} = \frac{k_{\mathrm{on}}\Gamma\alpha\bar{c} - (k_{\mathrm{off}} - k_{\mathrm{on}}\Gamma\beta)\Gamma_{\mathrm{captured}}}{\bar{U}} \quad \text{(S21)}$$

$$\frac{d\bar{c}}{dz} = -\frac{2}{R_{\mathrm{BAT}}}\frac{d\Gamma_{\mathrm{captured}}}{dz} \quad \text{(S22)}$$

Initial conditions are given as $\bar{c}(z = 0) = c_{\mathrm{in}}$, and $\Gamma_{\mathrm{captured}}(z = 0) = 0$. By integrating Eq. S22 and considering the initial conditions,

$$\Gamma_{\mathrm{captured}} = \frac{R_{\mathrm{BAT}}}{2}(c_{\mathrm{in}} - \bar{c}) \quad \text{(S23)}$$

By inserting Eqs. S22 and S23 into Eq. S21,

$$\frac{d\bar{c}}{dz} = \frac{-(2\alpha k_{\mathrm{on}}\Gamma/R_{\mathrm{BAT}} + k_{\mathrm{off}} - k_{\mathrm{on}}\Gamma\beta)\bar{c} + (k_{\mathrm{off}} - k_{\mathrm{on}}\Gamma\beta)c_{\mathrm{in}}}{\bar{U}} \quad \text{(S24)}$$

Eq. S24 is separable as follows.

$$\int \frac{1}{-(2\alpha k_{\mathrm{on}}\Gamma/R_{\mathrm{BAT}} + k_{\mathrm{off}} - k_{\mathrm{on}}\Gamma\beta)\bar{c} + (k_{\mathrm{off}} - k_{\mathrm{on}}\Gamma\beta)c_{\mathrm{in}}}\, d\bar{c} = \int \frac{1}{\bar{U}}\, dz \quad \text{(S25)}$$

By integrating both sides of Eq. S25 and rearranging the equation, we obtain

$$\bar{c}(z) = \frac{(k_{\mathrm{off}} - k_{\mathrm{on}}\Gamma\beta)c_{\mathrm{in}}}{2\alpha k_{\mathrm{on}}\Gamma/R_{\mathrm{BAT}} + k_{\mathrm{off}} - k_{\mathrm{on}}\Gamma\beta}\left(1 + Ae^{\frac{-(2\alpha k_{\mathrm{on}}\Gamma/R_{\mathrm{BAT}} + k_{\mathrm{off}} - k_{\mathrm{on}}\Gamma\beta)z}{\bar{U}}}\right) \quad \text{(S26)}$$

where $A$ is an arbitrary integration constant. Taking the initial condition, $\bar{c}(0) = c_{\mathrm{in}}$,

$$\bar{c}(z)=\frac{(k_{\mathrm{off}}-k_{\mathrm{on}}\Gamma\beta)c_{\mathrm{in}}}{2\alpha k_{\mathrm{on}}\Gamma/R_{\mathrm{BAT}}+k_{\mathrm{off}}-k_{\mathrm{on}}\Gamma\beta}\left(1+\frac{2\alpha k_{\mathrm{on}}\Gamma/R_{\mathrm{BAT}}}{k_{\mathrm{off}}-k_{\mathrm{on}}\Gamma\beta}e^{\frac{-(2\alpha k_{\mathrm{on}}\Gamma/R_{\mathrm{BAT}}+k_{\mathrm{off}}-k_{\mathrm{on}}\Gamma\beta)z}{\bar{U}}}\right) \quad \text{(S27)}$$

The control slice leaves the BAT at $z = L_{\mathrm{BAT}}$, therefore the average concentration at the outlet, $c_{\mathrm{out}}$ is

$$c_{\mathrm{out}}=\frac{(k_{\mathrm{off}}-k_{\mathrm{on}}\Gamma\beta)c_{\mathrm{in}}}{2\alpha k_{\mathrm{on}}\Gamma/R_{\mathrm{BAT}}+k_{\mathrm{off}}-k_{\mathrm{on}}\Gamma\beta}\left(1+\frac{2\alpha k_{\mathrm{on}}\Gamma/R_{\mathrm{BAT}}}{k_{\mathrm{off}}-k_{\mathrm{on}}\Gamma\beta}e^{\frac{-(2\alpha k_{\mathrm{on}}\Gamma/R_{\mathrm{BAT}}+k_{\mathrm{off}}-k_{\mathrm{on}}\Gamma\beta)L_{\mathrm{BAT}}}{\bar{U}}}\right) \quad \text{(S28)}$$

BAT efficiency $\eta$ is defined as

$$\eta=1-\frac{c_{\mathrm{out}}}{c_{\mathrm{in}}}=\frac{2\alpha k_{\mathrm{on}}\Gamma/R_{\mathrm{BAT}}}{2\alpha k_{\mathrm{on}}\Gamma/R_{\mathrm{BAT}}+k_{\mathrm{off}}-k_{\mathrm{on}}\Gamma\beta}\left(1-e^{\frac{-(2\alpha k_{\mathrm{on}}\Gamma/R_{\mathrm{BAT}}+k_{\mathrm{off}}-k_{\mathrm{on}}\Gamma\beta)L_{\mathrm{BAT}}}{\bar{U}}}\right) \quad \text{(S29)}$$

Substituting back for $\alpha$ and $\beta$ defined in Eq. S15,

$$\eta=\frac{2k_{\mathrm{on}}\Gamma/R_{\mathrm{BAT}}}{2k_{\mathrm{on}}\Gamma/R_{\mathrm{BAT}}+k_{\mathrm{off}}}\left(1-e^{\frac{-(2k_{\mathrm{on}}\Gamma/R_{\mathrm{BAT}}+k_{\mathrm{off}})L_{\mathrm{BAT}}}{\bar{U}(1+\frac{k_{\mathrm{on}}\Gamma}{k_c})}}\right) \quad \text{(S30)}$$

From Eq. S7, the surface Damköhler number is defined as $\mathrm{Da_s} = k_{\mathrm{on}}\Gamma/k_{\mathrm{c}}$, and we define the effective surface on-rate $\kappa_{\mathrm{on}} = 2k_{\mathrm{on}}\Gamma/R_{\mathrm{BAT}}$. Thus, we get

$$\eta=\frac{\kappa_{\mathrm{on}}}{\kappa_{\mathrm{on}}+k_{\mathrm{off}}}\left(1-e^{\frac{-(\kappa_{\mathrm{on}}+k_{\mathrm{off}})L_{\mathrm{BAT}}}{(1+\mathrm{Da_s})\bar{U}}}\right) \quad \text{(S31)}$$

In this study, $\kappa_{\mathrm{on}}$ (~$10^{-1}$–$10^{2}$ $\mathrm{s^{-1}}$) >> $k_{\mathrm{off}}$ (~$10^{-5}$ $\mathrm{s^{-1}}$). Therefore,

$$\eta=1-e^{\frac{1}{1+\mathrm{Da_s}}\cdot\frac{-\kappa_{\mathrm{on}}L_{\mathrm{BAT}}}{\bar{U}}} \quad \text{(S32)}$$

Because from Eq. S5, the first Damkhöler number is defined as $\mathrm{Da_I} = \kappa_{\mathrm{on}}L_{\mathrm{BAT}}/\bar{U}$, we can rewrite Eq. S32 as:

$$\eta=1-e^{\frac{-\mathrm{Da_I}}{1+\mathrm{Da_s}}} \quad \text{(S33)}$$

For the model analyte with $D$ = 80 μm$^2$ s$^{-1}$ flowing through a BAT with $R_{\mathrm{BAT}}$ = 4 μm and $L_{\mathrm{BAT}}$ = 20 μm, the surface Damköhler number, $\mathrm{Da_s} < 1$ under the experimental conditions in this study (Fig. S1b,c). Therefore, for the experimental conditions used in this study, Eq. S33 simplifies to:

$$\eta=1-e^{-\mathrm{Da_I}} \quad \text{(S34)}$$

The BAT efficiency $\eta$ was found to be consistent with FEM, SBAD and experimental results as shown in Fig. 3d,e.

**Supplementary Discussion S3. Stochastic Brownian agent dynamics (SBAD)**

In this section, we discuss key design considerations and the algorithm of the stochastic Brownian agent dynamics (SBAD) analysis. SBAD is an agent-based method that models and traces an analyte flowing through a BAT by superposing stochastic random walk (i.e. Brownian motion) with the displacement by convective flow, detects stochastic wall collision and further models binding as a random, stochastic event that only occurs for a subset of collisions, Fig. S3. The probability of affinity binding upon wall collision is defined as $P_{ab}$. $P_{ab} = 1$ in a scenario where every collision would lead to binding. In our models $P_{ab} << 1$. To model the stochasticity, a random number $0 \leq$ number $\leq 1$ is generated, and only if the number $\leq P_{ab}$ does binding occur.

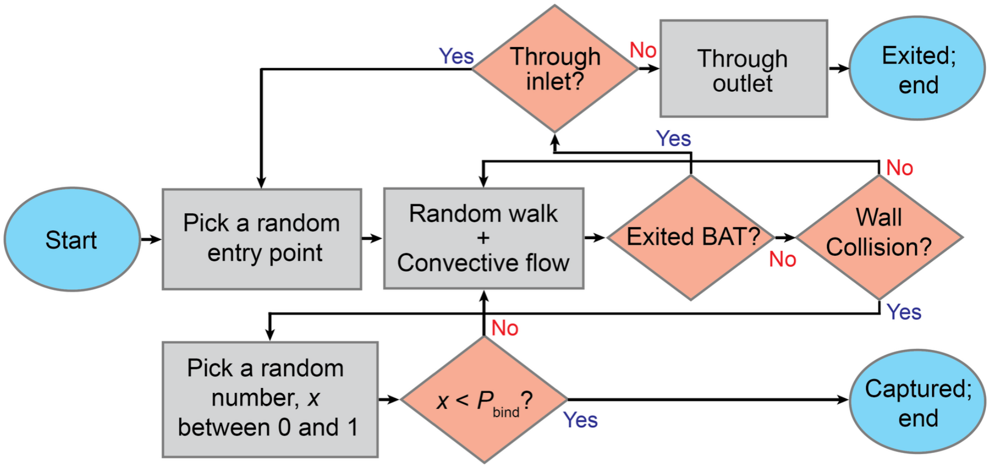


**Fig. S3 | Stochastic Brownian agent dynamics (SBAD) algorithm for flow, random walk, wall collision and capture of an analyte entering a BAT.** The SBAD algorithm was programmed and executed using MATLAB.

In SBAD, the circular, axisymmetric BAT is represented as a 2D microchannel. An analyte enters the BAT at a randomly location of the inlet, and at every time step ($\Delta t$ = 10 μs), SBAD updates the analyte location by superposing convective transport assuming a parabolic flow profile neglecting the entrance length ($L_{entrance} << L_{BAT}$), and a random walk representing Brownian motion. From diffusion equation[8], the displacement of analytes undergoing Brownian motion in 2D is expressed as a normal distribution with zero mean (initial point) and variance of $4Dt$ at time $t$. Thus, at each time step $\Delta t$, while randomly drawing the displacement direction from all possible directions (0–2π rad) to reflect the isotropic nature of Brownian motion, we sample the displacement from half-normal distribution of zero mean and variance of $4D$t (because the direction is already defined). In the event that the analyte leaves through the exit ($z \geq L_{BAT}$), the simulation ends, and the analyte is recorded as exited. Conversely, if the analyte leaves through the entrance ($z \leq 0$), the simulation is reset because the analyte is assumed to re-enter a BAT through a new random entry point. Once the collision of an analyte with the BAT wall is detected, the analyte is affinity bound if the random number $\leq P_{ab}$, or if not, the random walk simulation continues from the wall position.

Calibration of SBAD with FEM models.

SBAD uses Monte Carlo estimates of repeated simulations to calculate distributions and capture efficiency $\eta$. To calibrate SBAD, the free parameter $P_{ab}$ was determined by benchmarking $\eta$ against deterministic ensemble FEM results. We calibrated $P_{ab}$ as a function of $\kappa_{on}$ and $D$.

Since each analyte flowing through a BAT is either captured or escapes, statistical SBAD estimates for $\eta$ follow a Bernoulli distribution with mean $\eta$ and variance $\eta\,(1-\eta)$. A SBAD analysis of $N$ analytes helps establish $\eta$ with a standard error SE:

$$\mathrm{SE} = \sqrt{\frac{\eta(1-\eta)}{N}} \tag{S35}$$

To ensure a precise calibration of $P_{ab}$, we statistically choose the number of analytes $N$ that gives a two-sided 95% confidence interval (CI) which is 1.96 SE to be no wider than ±5% of the calculated value.

$$1.96\mathrm{SE} < 0.05\eta \tag{S36}$$

$$N > \frac{1.96^2(1-\eta)}{0.05^2\eta} \approx 1{,}536\frac{1-\eta}{\eta} \tag{S37}$$

With the minimum requirement of $N$ = 5,000 and by selecting the first multiple of 1,000 that satisfies the constraint, we find the values of $P_{ab}$ that yield the 95% CIs of $\eta$ matching the FEM benchmark data. For $0.235 < \eta < 1$, we ran 5,000 simulations, for $0.204 < \eta < 0.235$, 6,000 simulations, for $0.180 < \eta < 0.204$, 7,000 simulations, for $0.161 < \eta < 0.180$, 8,000 simulations and so on. At $\bar{U} = 10\ \mu\mathrm{m\ s^{-1}}$, $P_{ab}$ was calibrated across $\kappa_{on}$ for a 100 kDa model globular protein analyte with $D = 80\ \mu\mathrm{m^2\ s^{-1}}$ and across $D$ at $\kappa_{on} = 5\ \mathrm{s^{-1}}$ in the same order of magnitude that can be obtained from typical capture surfaces in this study, Extended Data Fig. 2.

**Supplementary Discussion S4. Selective functionalization of the BATs in the MFSD by float-flip-float incubation**

The MFSD is premised on exclusive binding of analytes inside of the BATs following Brownian-motion induced wall collision. BATs are partitioned by immersion into a fluorinated oil which displaces water from the intrinsically hydrophobic MFSD membrane, while trapping the aqueous solution in the BATs. The BATs must, however, be functionalized with capture probes that will bind analytes with high affinity. The hydrophobic MFSD was functionalized by N-Hydroxysuccinimide (NHS)-chemistry via UV-induced grafting of the NHS ester through residual unreacted methacrylate groups on the membrane, subsequently immobilizing capture probes by substituting the NHS ester on the surface.

As a conventional surface treatment approach, one might think of submerging the MFSD in an activation solution, followed by chemical immobilization of the capture probe. However, this process functionalizes not only the inside of the BAT but also the top and bottom surfaces of the MFSD, Extended Data Fig. 3a. Consequently, the top and bottom surfaces become hydrophilic and capable of capturing analytes. This causes ineffective partitioning of BATs, leaving a water film that leads to diffusive leakage across different BATs, Extended Data Fig. 3b. As a result, digital readout is not feasible because the number of true positive BATs cannot be reliably determined.

We introduce a float-flip-float incubation for selective functionalization of the inside of the BAT pores. The MFSD membrane was not submerged but floated by surface tension on the solution containing acrylic acid-NHS-ester, with each pore being filled and acting as a stop valve preventing the liquid from reaching the 'top surface'. Hence only the 'bottom surface' and the inside of the BATs were exposed to the solution, and upon UV exposure, were functionalized with reactive NHS-ester, while the 'top surface' remained pristine. The MFSD membrane was then washed, flipped, and floated on a solution containing capture probe molecules with amine groups. Again, the BATs were filled with the solution by capillary flow formed stop valves, and probes reacted exclusively with the previously activated surface which was only inside of the BATs. Next, the membrane was rinsed, and exposed to a solution containing 1H,1H-heptafluorobutylamine (*i.e.* fluorocarbon-amine) to quench unreacted NHS-ester on the 'bottom surface', and restore the double-sided hydrophobicity, Extended Data Fig. 3c. Digital readout test using an MFSD functionalized with the float-flip-float method shows effective confinement of the fluorescent signals within individual BATs, thus supporting a digital readout, Extended Data Fig. 3d.

**Supplementary Discussion S5. Varying and measuring $\kappa_{on}$**

In this study, $\kappa_{on}$ was varied from 0.0744 to 74.4 $s^{-1}$ by using five different capture-analyte systems including (i) biotin-SβG, (ii&iii) antibody-antigen (anti-streptavidin and anti-βG antibodies for SβG), and a nucleic acid capture probe against biotinylated oligonucleotide analytes with (iv) complementary and (v) double-mismatch sequences, both pre-coupled with SβG (Fig. 3e). To estimate $\kappa_{on}$, we performed ensemble measurement of $k_{on}$ and $\Gamma$. Despite the imprecision of a coefficient of variation (CV) on the order of tens of percents for each measurement, experimental results with MFSDs and single-molecule counting showed strong consistency with analytical and FEM predictions as well as SBAD when using the experimentally estimated $\kappa_{on}$. This agreement validates the experimentally estimated $\kappa_{on}$.

The on-rate $k_{on}$ of capture probes to their analytes were measured using SPR for the antibodies (Fig. S4) and obtained from literatures for biotin and nucleic acids[9,10]. The surface densities $\Gamma$ of capture probes were quantified using two different approaches depending on the probe affinities. For high-affinity probes such as biotin ($K_D \sim 10^{-14}$ M), and nucleic acids ($K_D \sim 10^{-13}$ M), reporter solutions were incubated with functionalized MFSDs and reporter depletion caused by surface binding was measured. In this case, we presume that the surfaces are saturated, and the experimental conditions are chosen such that a substantial depletion of the reporter solution occurs, and the extent of reporter depletion thus directly reflects probe density. The differences between reporter depletions by the pristine and functionalized MFSD were used to quantify the surface density $\Gamma$ (Fig. S5a–d). 4'-hydroxyazobenzene-2-carboxylic acid (HABA) and a fluorescently labelled complementary DNA oligonucleotide were used as a reporter molecule and $\Gamma$ of biotin and nucleic acid were measured as $4.96 \times 10^{-8}$ mol $m^{-2}$ (CV = 25%; $n$ = 3) and $3.65 \times 10^{-10}$ mol $m^{-2}$ (CV = 24%; $n$ = 3). The $\Gamma$ of biotin was then varied based on ratiometric incorporation of spacer molecules, wherein biotin was introduced as biotin-PEG-amine and the spacer as methyl-PEG-amine at NHS chemistry-based surface coating.

The measurement of antibody surface density coverage faced additional challenges because antibodies bind more weakly ($K_D \sim 10^{-11}$ M) and saturation on the surface cannot be reached except for very high concentrations. However, for high concentrations, the loss in reagents and depletion become too small to measure accurately. Instead, antibody densities were determined from the depletion of the surface coating solution itself, where the reduction in free capture probe concentration reflects immobilization onto the surface. The differences between free capture probe depletions with and without an MFSD were used to quantify the surface density. (Fig. S5e,f). To quantify the antibody depletion in the coating solution, a bicinchoninic acid (BCA) assay was performed for solutions incubated with and without an MFSD, and the calculated antibody densities were $2.08 \times 10^{-8}$ mol m$^{-2}$ (CV = 10%; $n$ = 3) and $1.25 \times 10^{-8}$ mol m$^{-2}$ (CV = 20%; $n$ = 3) for anti-streptavidin and anti-βG antibodies, respectively. The measured $k_{on}$, $\Gamma$, and $\kappa_{on}$ are summarized in Extended Data Table 1.

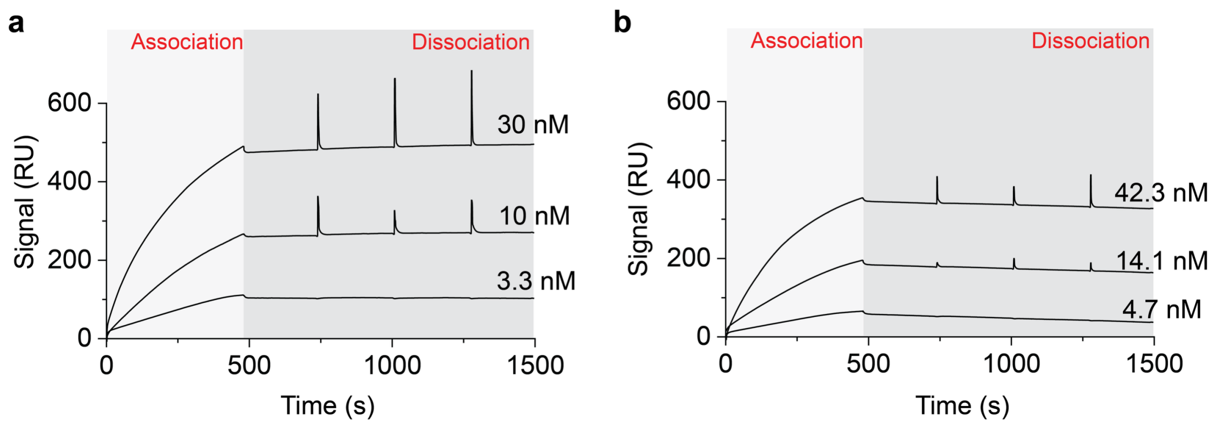


**Fig. S4 | Surface plasmon resonance (SPR) sensorgrams.** SPRs were run for both anti-streptavidin and anti-β-galactosidase antibodies with 480 s of incubation and association (light grey) followed by washing and 1000 s of dissociation (dark grey). On-rates and off-rates were obtained by fitting the sensorgrams. **a**, Sensorgram for anti-streptavidin antibody. The on-rate and off-rate were evaluated as $k_{on} = 4.55 \times 10^2$ m$^3$ mol$^{-1}$ s$^{-1}$ and $k_{off} = 7.01 \times 10^{-6}$ s$^{-1}$. **b**, Sensorgram for anti-β-galactosidase antibody. The on-rate and off-rate were evaluated as $k_{on} = 3.65 \times 10^2$ m$^3$ mol$^{-1}$ s$^{-1}$ and $k_{off} = 2.01 \times 10^{-4}$ s$^{-1}$. The spikes in the dissociation phase (dark grey) were caused by syringe switching in the SPR system.

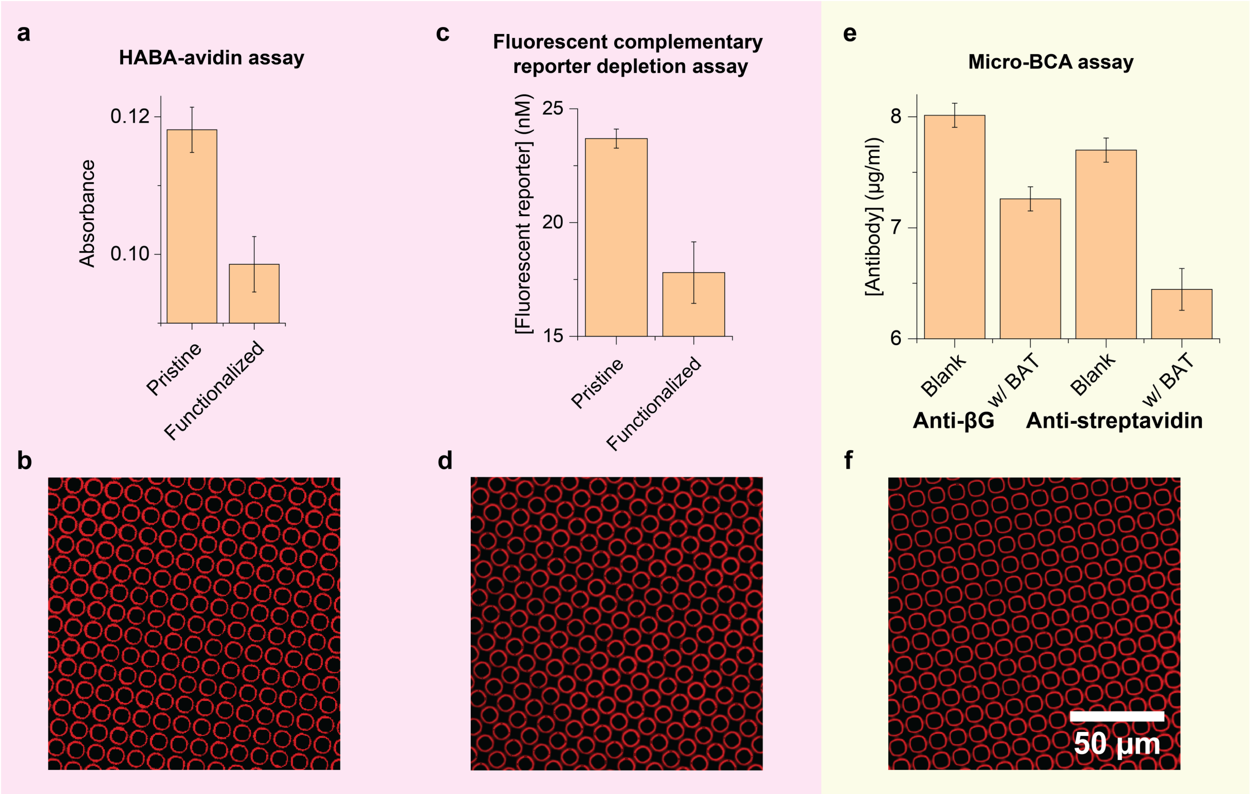


**Fig. S5 | Surface density measurements.** The surface density was measured by either reporter depletion for biotin and nucleic acid (red shading) or free capture probe depletion in the coating solution for antibodies (yellow shading). **a**, Surface density measurement of biotin. 4'-hydroxyazobenzene-2-carboxylic acid (HABA)-avidin assay with absorbance at 500 nm were performed after incubation with pristine and functionalized MFSDs. **b**, Fluorescently labelled biotin-coated MFSD. Red fluorescence visualizes surface-immobilized biotin. **c**, Surface density measurement of oligonucleotide capture probe. Concentrations of fluorescent complementary DNA oligonucleotide reporter were measured after incubation with pristine and functionalized MFSDs. **d**, Fluorescently labelled oligonucleotide-coated MFSD. Red fluorescence visualizes surface-immobilized oligonucleotide probe. **e**, Surface density measurement of antibodies. Coating solution concentrations were measured after incubating with and without an MFSD by the bicinchoninic acid (BCA) assay. **f**, Fluorescently labelled antibody-coated MFSD. Red fluorescence visualizes surface-immobilized antibody.

**Supplementary Discussion S6. Determination of the zeptomolar LOD from Poisson and experimental variability.**

At zM concentrations, only a few analyte molecules are expected in the 200 μL sample, and therefore discrete molecular sampling and low-count background variability become important in defining the LOD. For the one-step MFSD test for SβG detection, we consider both the Poisson and the experimental variation across blank measurements. In the one-step MFSD test, no SβG was present in the blank and therefore no positive BATs were observed as expected. However, the dynamic thresholding used to classify BATs as positive or negative produced a small false-positive fraction. The measured blank signal was $3.4 \times 10^{-4}$ AEB on average, corresponding to approximately 170 BATs classified as positive among 500,000 BATs.

The experimental detection threshold was defined as the mean blank AEB plus three standard deviations measured from blank samples, yielding $6.4 \times 10^{-4}$ AEB. The fitted standard curve intersects this threshold at 215 zM, corresponding to an expected ~26 molecules in a 200 μL sample.

We additionally consider the Poisson variation associated with the ~170 background false positive BATs where the standard deviation is obtained as $\sqrt{170} \approx 13.0$ BATs, giving a three times the standard deviation of 39 counts, which the number of captured analyte molecules should exceed for detection above the background. Accounting for the measured 50% capture efficiency, this corresponds to 78 molecules in the 200 μL sample or 648 zM.

We therefore report both 215 zM as the experimental LOD and 648 zM as a Poisson LOD for robust low-copy detection. Importantly, both are in the zM regime: the lower experimental LOD reflects the sensitivity reached in this experimental calibration, whereas the Poisson LOD provides a conservative estimate that accounts for stochastic low-copy variation.

**Table S1 | MFSD mass transport and analyte capture parameters for SβG, IL-4 and Influenza A nucleoprotein.** Calculations were performed for BATs with $R_{BAT}$ = 4 μm and $L_{BAT}$ = 20 μm, at the sample-flow velocity $\bar{U}$ = 133 μm $s^{-1}$. For SβG detection, BATs were coated with biotin at $\Gamma = 0.496 \times 10^{-8}$ mol $m^{-2}$ (See Extended Data Table 1). For antibody-functionalized BATs, a surface density of $\Gamma = 2 \times 10^{-8}$ mol $m^{-2}$ was used based on the experimentally measured antibody surface density obtained using the same surface functionalization method.

| Parameter | SβG | IL-4 | Influenza A nucleoprotein | Basis |
|---|---|---|---|---|
| Molecular weight, $M$ (kDa) | 530 | 15 | 56.7 | SβG: Experimental validation<br>IL-4 & Influenza A nucleoprotein: vendor specification |
| Molecular diameter, $d_{molecule}$ (nm) | 10.7 | 3.27 | 5.09 | $d_{molecule} = \left(\frac{6M}{\pi \rho N_A}\right)^{\frac{1}{3}}$<br>$\rho$: Density of protein, for globular proteins, 1.37 g $cm^{-3}$<br>$N_A$: Avogadro's number, $6.02 \times 10^{23}$ $mol^{-1}$ |
| $D$ ($\mu m^2\ s^{-1}$) | 45.9 | 150 | 96.4 | $D = \frac{k_B T}{3\pi\mu d_{molecule}}$<br>$k_B$: Boltzmann constant, $1.381 \times 10^{-23}$ J $K^{-1}$<br>$T$: Temperature, 298 K (25 ºC)<br>$\mu$: Dynamic viscosity, for water at 25 ºC, $0.89 \times 10^{-23}$ N s $m^{-2}$ |
| Gz | 9.27 | 2.84 | 4.41 | Eq. S4 |
| Sh | 4.19 | 3.84 | 3.93 | Eq. S10 |
| $k_c$ ($\mu m\ s^{-1}$) | 24.0 | 72.0 | 47.4 | Eq. S9 |
| $k_{on}$ ($m^3\ mol^{-1}\ s^{-1}$) | $3.00 \times 10^3$ | $5.57 \times 10^3$ | $2.52 \times 10^1$ | Literature reported[10-12] |
| $Da_s$ | 0.620 | 1.55 | 0.0106 | Eq. S7 |
| $\kappa_{on}$ ($s^{-1}$) | 7.44 | 55.7 | 0.252 | $\kappa_{on} = \frac{2k_{on}\Gamma}{R_{BAT}}$ |
| $\tau_{res}$ (s) | 0.150 | 0.150 | 0.150 | $\tau_{res} = \frac{L_{BAT}}{\bar{U}}$ |
| $Da_I$ | 1.12 | 8.36 | 0.0378 | Eq. S5 |
| $\eta$ | 49.9% | 96.2% | 3.67% | Eqs. 1 and 2 |

#### Supplementary Videos

**Video S1. Stochastic Brownian agent dynamics (SBAD) simulation for 10 analytes entering a Brownian affinity trap (BAT) with 50% capture efficiency.** The BAT has a 4-μm radius and is 20-μm long, and coated with a capture probe. It has an effective on-rate $\kappa_{on}$ = 5 $s^{-1}$ and a corresponding affinity binding probability $P_{ab}$ = 0.014. The analytes have a diffusion coefficient of $D$ = 80 $\mu m^2 s^{-1}$ and flow at an average flow speed of $\bar{U}$ = 133 $\mu m\ s^{-1}$, which yields an 50% capture efficiency. The SBAD simulations were run independently for each of the 10 analytes with a time step of $10^{-5}$ s, and simulation results were rendered as a slow-motion animation showing all analytes flowing simultaneously at a playback speed of 0.01×.

**Video S2. SBAD simulation for 10 analytes entering three BATs with different flow speeds $\bar{U}$.** 10 analytes with $D$ = 80 $\mu m^2 s^{-1}$ flow with different average flow speeds of $\bar{U}$ = 50, 133 and 1190 $\mu m\ s^{-1}$, yielding a 70, 50 and 10% capture efficiency, respectively. As previously, each of the three BATs has a radius of 4 μm and a length of 20 μm, and coated with a capture probe at $\kappa_{on}$ = 5 $s^{-1}$ which corresponds to $P_{ab}$ = 0.014. The SBAD simulations were run independently for each analyte with the time step of $10^{-5}$ s, and simulation results were rendered as a slow-motion animation showing all analytes flowing simultaneously at a playback speed of 0.01×.

**Video S3. SBAD simulation for 10 analytes entering three BATs with different binding probabilities $P_{ab}$ and the corresponding effective on-rates $\kappa_{on}$.** The analytes flow through three BATs each coated with a capture probe with different $\kappa_{on}$ = 80, 5 and 0.7 $s^{-1}$, which correspond to $P_{ab}$ = 0.36, 0.014 and 0.0014, yielding a 70, 50 and 10% capture efficiency, respectively. Again, the three BATs had a radius of 4 μm and a length of 20 μm and the analytes were flowed at $\bar{U}$ = 133 $\mu m\ s^{-1}$ and their $D$ = 80 $\mu m^2 s^{-1}$. The SBAD simulations were run independently for each analyte with the time step of $10^{-5}$ s, and simulation results were rendered as a slow-motion animation showing all analytes flowing simultaneously at a playback speed of 0.01×.

#### References

1 Bird, R. B., Stewart, W. & Lightfoot, E. N. *Transport Phenomena second edition*. (John Wiley & Sons, 2002).

2 Gervais, T. & Jensen, K. F. Mass transport and surface reactions in microfluidic systems. *Chem. Eng. Sci.* **61**, 1102-1121 (2006).

3 Squires, T. M., Messinger, R. J. & Manalis, S. R. Making it stick: convection, reaction and diffusion in surface-based biosensors. *Nat. Biotechnol.* **26**, 417-426 (2008).

4 Fawzy, M. K., Varela-Corredor, F., Boi, C. & Bandini, S. The Role of the Morphological Characterization of Multilayer Hydrophobized Ceramic Membranes on the Prediction of Sweeping Gas Membrane Distillation Performances. *Membranes* **12**, 939 (2022).

5 Deen, W. M. *Analysis of Transport Phenomena*. (OUP USA, 1998).

6 Brown, G. M. Heat or mass transfer in a fluid in laminar flow in a circular or flat conduit. *AIChE Journal* **6**, 179-183 (1960).

7 White, F. M. *Fluid mechanics*. (McGraw-Hill, 2015).

8 Risken, H. in *The Fokker-Planck equation: methods of solution and applications* (Springer, 1989).

9 Todisco, M., Ding, D. & Szostak, J. W. Transient states during the annealing of mismatched and bulged oligonucleotides. *Nucleic Acids Res.* **52**, 2174-2187 (2024).

10 Srisa-Art, M., Dyson, E. C., deMello, A. J. & Edel, J. B. Monitoring of Real-Time Streptavidin−Biotin Binding Kinetics Using Droplet Microfluidics. *Anal. Chem.* **80**, 7063-7067 (2008).

11 Leirs, K. *et al.* Bioassay Development for Ultrasensitive Detection of Influenza A Nucleoprotein Using Digital ELISA. *Anal. Chem.* **88**, 8450-8458 (2016).

12 Rao, E., Mikol, V., Li, D., Kruip, J. & Davison, M. Antibodies that bind IL-4 and/or IL-13 and their uses. United States patent (2020).
